# Modeling and inference of spatial intercellular communications and multilayer signaling regulations using stMLnet

**DOI:** 10.1101/2022.06.27.497696

**Authors:** Jinyu Cheng, Lulu Yan, Qing Nie, Xiaoqiang Sun

## Abstract

Multicellular organisms require intercellular and intracellular signaling to coordinately regulate different cell functions. Although many methods of cell-cell communication (CCC) inference have been developed, they seldom account for both the intracellular signaling responses and global spatial information. The recent advancement of spatial transcriptomics (ST) provides unprecedented opportunities to better decipher CCC signaling and functioning. In this paper, we propose an <u>ST</u>-based <u>m</u>ultilayer <u>net</u>work method, stMLnet, for inferring spatial intercellular communication and multilayer signaling regulations by quantifying distance-weighted ligand–receptor signaling activity based on diffusion and mass action models and mapping it to intracellular targets. We benchmark stMLnet with existing methods using simulation data and 8 real datasets of cell type-specific perturbations. Furthermore, we demonstrate the applicability of stMLnet on six ST datasets acquired with four different technologies (e.g., seqFISH+, Slide-seq v2, MERFIS and Visium), showing its effectiveness and reliability on ST data with varying spatial resolutions and gene coverages. Finally, stMLnet identifies positive feedback circuits between alveolar epithelial cells, macrophages, and monocytes via multilayer signaling pathways within a COVID-19 microenvironment. Our proposed method provides an effective tool for predicting multilayer signaling regulations between interacting cells, which can advance the mechanistic and functional understanding of spatial CCCs.

## Introduction

It has become increasingly clear that cell fates and functions are not only determined by the intrinsic genetic makeup of the cell, but they are also influenced by neighboring cells and the spatial environment in multicellular tissue ^1,2^. Recent studies have revealed that cell–cell interactions (CCIs) play important roles in cell differentiation, tissue development, immunity, and cancers ^3^. Elucidating the mechanisms by which intercellular signaling regulates intracellular gene expression is essential to advance our understanding of the functional and therapeutic roles of CCIs ^4^.

Cells can interact with each other in multiple ways, such as physical cell–cell contact (e.g., cell adhesions) ^5,6^ and biochemical cell–cell communication (CCC) mediated by diffusible molecules (e.g., autocrine, paracrine or endocrine via ligand–receptor (LR) interactions) ^4^. CCC plays vital roles in many physiological functions of multicellularity, and its dysregulation often drives the occurrence and development of many diseases ^7^. CCC generally involves four sequential events ^8^: 1) secretion and diffusion of ligand molecules; 2) reception and specific binding of the ligand by the receptor, leading to receptor activation; 3) induction of one or multiple signaling transduction pathways by the activated receptors, leading to the activation of downstream transcription factors (TFs); and 4) regulation of target gene (TG) expression by the activated TFs and initiation of cell phenotype switching. Together, the signaling mechanisms involved in functional CCC include both intercellular LR interactions and intracellular signaling transduction and transcriptional regulation, which is referred to as the multilayer signaling network ^9-11^. Traditional experimental studies have often focused on a few signaling molecules to study intercellular communications, lacking information across multiple scales of signaling. Systems analysis of signaling regulations in functional CCC is challenging and critical.

Single-cell RNA sequencing (scRNA-seq) provides unprecedented high-throughput data to decipher intercellular communication by analyzing cell-type-specific expression of ligands and receptors ^4^. Recently, several computational methods or tools have been developed to infer CCC from scRNA-seq data ^4,11^. However, most existing methods or tools (e.g., ^12-15^) only consider intercellular LR signaling pairs without dissecting the response of their downstream pathways (see a comprehensive comparison of related methods in **Table S1**).

Previously we developed a multilayer network approach to infer both intercellular and intracellular signaling networks ^9,16,17^. While several other methods (e.g., ^18,19^) also consider intracellular responses, the information of cellular distance affecting the transportation and reception of secretory molecules across the spatial environment is missing because of the loss of the positional information of cells in the scRNA-seq data. Recently, the rapid advancement of spatial transcriptomics (ST) that measures both gene expression and positional information of cells ^20^ has afforded new opportunities to infer spatial and functional CCC ^21^.

In this study, we present stMLnet, an ST-based <u>m</u>ultilayer <u>net</u>work method for inferring spatial intercellular communication and intracellular signaling responses. stMLnet mechanistically quantifies the spatially dependent LR signaling activity based on a mathematical model of ligand diffusion and the LR interaction, and it quantitatively maps the intercellular LR signals to intracellular gene expressions using explainable tree-based machine learning. Our method can prioritize ligands, receptors, or their pairs that regulate specific target genes and can identify cellular circuits involving multilayer signaling regulations. We benchmark stMLnet with existing methods using simulation data and 8 real datasets of cell type-specific perturbations (e.g., those for breast cancer cell lines and glioma-bearing mice). Furthermore, we apply stMLnet to multiple ST datasets with varying spatial resolutions and gene coverages to show its effectiveness and robustness on ST data acquired with different technologies (e.g., seqFISH+, Slide-seq v2, and MERFIS as well as Visium). Moreover, we apply stMLnet to decipher intercellular and intracellular signaling mechanisms underlying immunotherapy resistance in gliomas and to analyze multilayer signaling feedback loops associated with inflammatory response to COVID-19 infection.

## Results

### Overview of stMLnet

Our proposed method aims to infer, quantify, and visualize CCCs in ST data with an emphasis on the multilayer signaling network and spatial cellular distance (**Fig 1**). First, stMLnet integrates multiple data sources of molecular interactions (including LR interaction, signaling pathways, and transcriptional regulations) into in-house knowledge databases (i.e., LigRecDB, RecTFDB, and TFTGDB) by using the algorithm of random walk with restart on directed weighted graphs (**Fig 1B**) (**Text S1-S2**). Second, based on the prior network information and gene expression data, stMLnet employs co-expression analysis and Fisher’s exact test to construct the structure of the multilayer network, acquiring the signaling paths from the upstream LR pairs to downstream target genes (LR∼TG) (**Fig 1C**). Third, to quantify LR signaling activity, stMLnet mechanistically models the LR signaling activity as a cell-distance-dependent function based on a ligand diffusion model and the law of mass action (**Fig 1D**). Lastly, to model the nonlinear LR-TG regulation relationships, stMLnet employs an explainable tree-based modeling approach (e.g., random forest regression) to link LR activity to target gene expression, with feature importance estimating the contribution of each upstream ligand and/or receptor to the target gene expression (**Fig 1E**).

**Fig 1.**
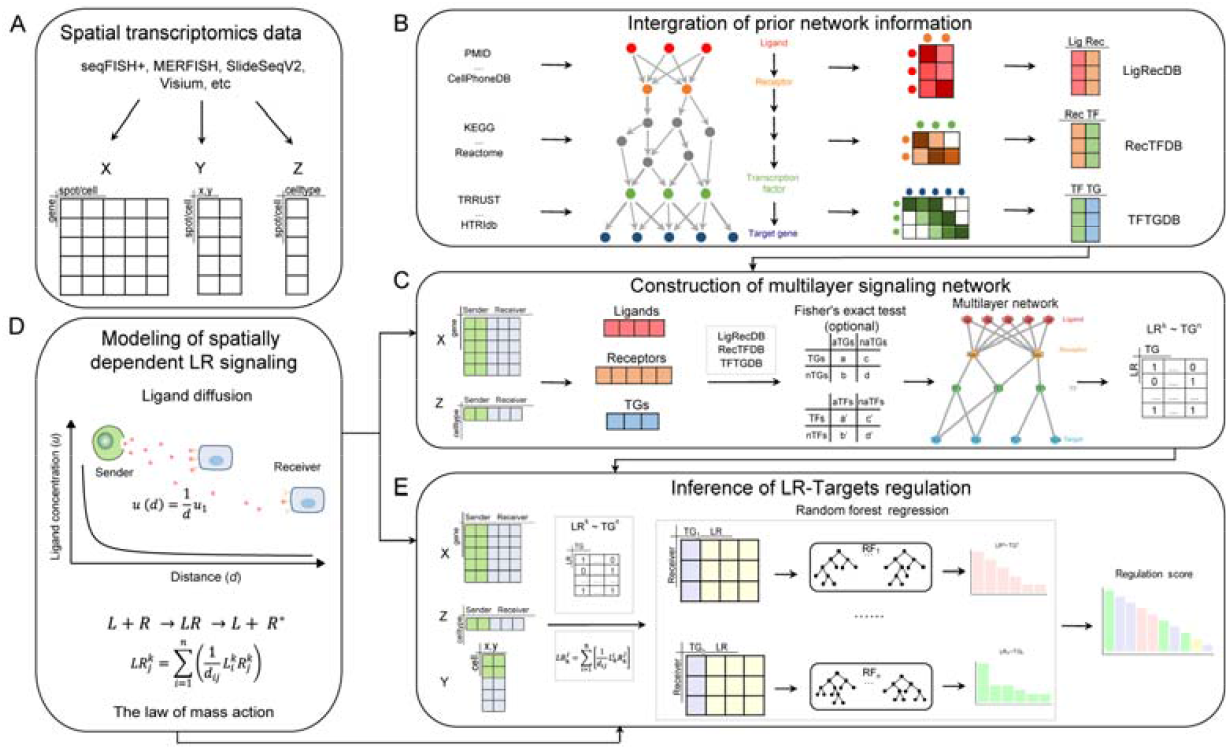
stMLnet framework. (**A**) Inputs of stMLnet, including the gene expression matrix (gene×cell), cell location matrix (cell×dim), and user-supplied cell type matrix (cell×type). (**B-E**) Workflow of stMLnet. (**B**) Integration of prior network information. stMLnet integrates prior information from multiple data sources of molecular interactions, including LR interaction, signaling pathways, and transcriptional regulations, into prior knowledge databases of stMLnet (i.e., LigRecDB, RecTFDB, and TFTGDB) based on directed weighted graphs and the random walk algorithm. (**C**) Statistical inference of multilayer signaling network. Combining the prior interaction knowledge and gene expression data, stMLnet infers multilayer signaling paths from the upstream LR pairs to downstream target genes (TGs) (LR∼TG). (**D**) Quantification of LR signaling activity. Based on the ligand diffusion model and the law of mass action, the LR signaling activity is modeled as a distance-dependent function. (**E**) Random forest regression links the LR activity to target gene expression. Explainable importance score of the specific LR signaling contributing to the target expression is computed to prioritize the upstream ligands or receptors.

The ***input*** of stMLnet includes the gene expression matrix (gene×cell) (***X***), cell location matrix (cell×dim) (***Y***), and user-supplied cell type matrix (cell×type) (***Z***). The cell location matrix is used to calculate the cell distance matrix (***D***). The cell type matrix indicates the pre-defined cell type annotation and labels of receiver cells and/or sender cells for CCC analysis (**Fig 2A**). For low-resolution ST data, such as Visium data, the users may use deconvolution methods to annotate the cell types in each spot, if needed ^22^. In addition, stMLnet also allows inputs of a set of genes of interest (e.g., the interaction-changed genes (ICGs) ^23^ or differentially expressed genes (DEGs)) as target genes for analysis. The ***output*** of stMLnet includes the multilayer signaling network (LR-TF-TG), LR signaling activity, and importance ranking of LR or L/R with respect to their abilities to regulate target gene expression. stMLnet also provides various ***visualizations*** of the results, for instance, the circle plot of the CCC network (**Fig 2B**), edge bundling plot of intercellular LR activity (**Fig 2C**), waterfall plot of multilayer signaling network (**Fig 2D**), multilayer network visualization of sub-networks downstream of a specified ligand (**Fig 2E**), and heatmap of functional enrichment for CCCs based on the enrichment analysis of downstream target genes (**Fig S1**).

**Fig 2.**
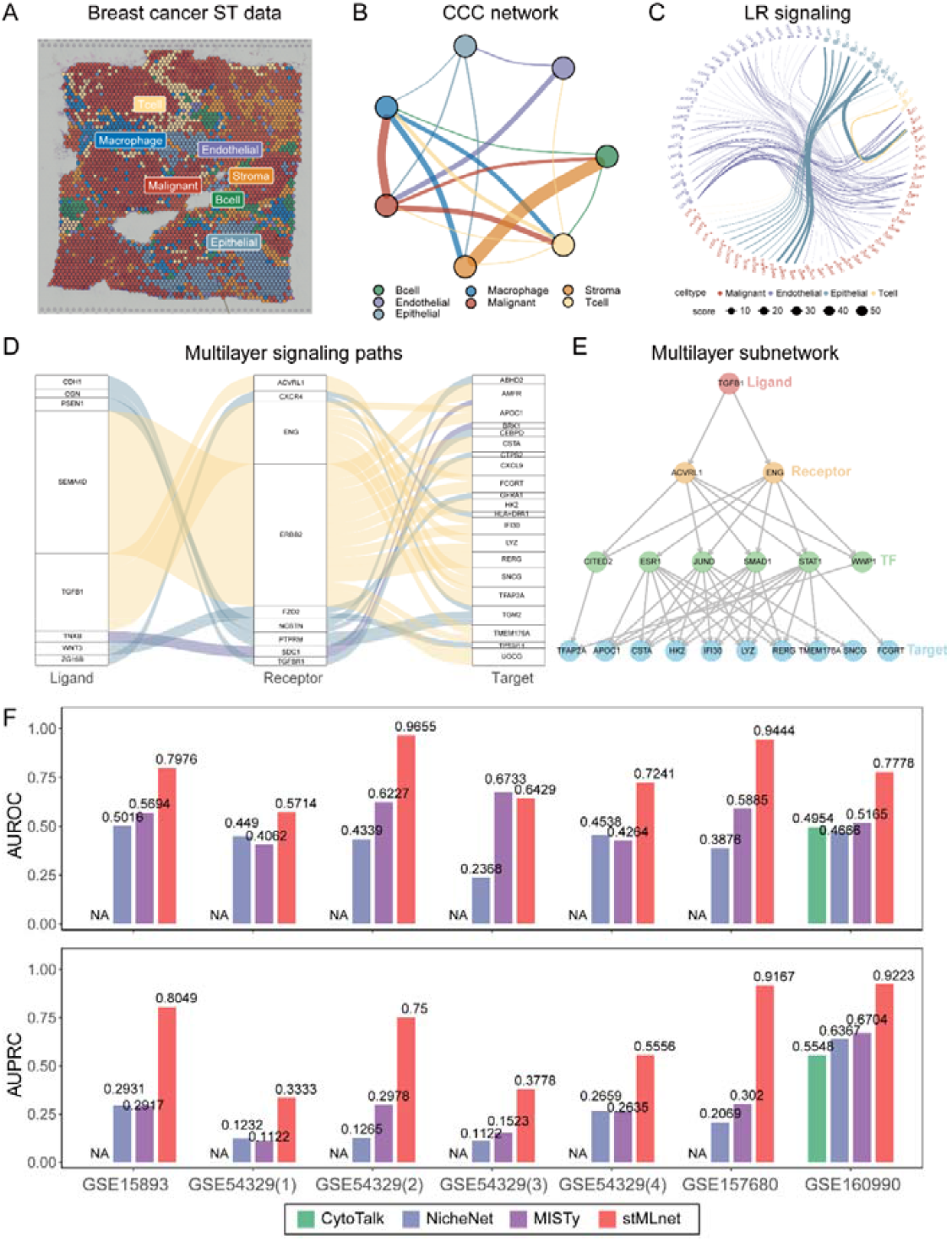
Validating stMLnet predictions using perturbation data. (**A**) The cell type annotation in the ST data of breast cancer. (**B**) The CCC network. Different node colors represent different cell types. The edges represent intercellular communications from the sender cells to the receiver cells. The edge color is consistent with that of the sender cells, and the edge width represents the number of LR pairs. (**C**) The edge bundling plot of LR signaling activity (top ranked). The nodes represent ligands or receptors, with different colors for different cell types. The edges represent LR signaling from the sender cells (e.g., T cells, endothelial cell, and epithelial cells) to the receiver cells (e.g., malignant cells here), with the edge color being consistent with that of the sender cell. The node size or edge width indicates the average strength of the LR signaling. (**D**) The waterfall plot of the multilayer signaling network. Regulatory paths from upstream LR pairs (top ranked) to their downstream targets in malignant cells are shown. The path color indicates the cellular source (sender cells) of the ligand signaling, and the path width represents the importance score of each LR with respect to TG regulation. (**E**) The multilayer signaling subnetwork indicates the regulatory paths from specified ligand/receptor(s) to the downstream TFs and then to target genes. Top-ranked target genes according to importance scores are prioritized for visualization. (**F**) Benchmarking stMLnet against other methods using cell line perturbation data. The prediction accuracy of stMLnet was compared to that of CytoTalk, NicheNet, or MISTy, assessed by AUCROC and AUCPR. ‘NA’ indicates that the corresponding ligands or receptors used for cell line perturbation could not be inferred by CytoTalk.

### Benchmarking stMLnet using perturbation data

To test the accuracy of the LR-target prediction of stMLnet on the breast cancer ST dataset ^24^, we collected 7 datasets of breast cancer cell lines with different perturbation conditions (e.g., TGFB1 or IL6 stimulation, NRP1 knockout, or CXCR4 mutation) (**Table S2**). If the downstream target genes are differentially expressed, they are considered to be potentially regulated by the corresponding ligand or receptor. As such, it is reasonable to use the DEGs as the ground truth of downstream targets of a specific ligand or receptor ^18,19^. To this end, we compared the predicted partial importance scores of the corresponding ligand or receptor on the target genes (Equation (2) in Methods section) with the ground truth (i.e., the differential expression status (true or false) of the target genes in each cell line after perturbation) and evaluated the prediction accuracy using AUROC (area under the ROC curve) and AUPRC (area under the Precision/Recall curve). The result (**Fig 2F**) indicates that stMLnet had good predictive accuracy on most datasets (AUROC was greater than 0.72, and AUPRC was greater than 0.55 on 5 of 7 datasets).

We also compared stMLnet with other methods of CCC inference or L/R-target regulation prediction. **Table S1** lists their characteristics, such as model/algorithm, input, intercellular prediction, intracellular prediction, and spatial information utilization. Among them, NicheNet ^18^, CytoTalk ^19^, and MISTy ^25^ were selected for quantitative benchmarking, as these three methods could predict L/R-target regulation. The implementation of these methods is described in **Text S3**. The benchmarking result (**Fig 2F**) indicates that stMLnet significantly outperformed the other three methods on all the 7 datasets.

To further evaluate the predictions of stMLnet regarding LR-target regulations, we hypothesized that the more reliable the inferred network, the higher the correlation between LR signaling activity and target gene expression. As such, we computed the mutual information (MI) or the Pearson correlation coefficient (PCC) for each pair of 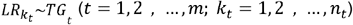 in the multilayer networks inferred by stMLnet based on the breast cancer ST dataset. We then compared values of MI or PCC of 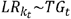 with that of random pairing or that inferred by other methods. Here, only CytoTalk could be used for comparison, as NicheNet predicts ligand-target regulations but does not involve receptors, and MISTy does not explicitly model LR interactions. The LR-target pairings predicted by stMLnet had larger MI values than CytoTalk and random pairing, across various cell types (**Fig S2AB**). In addition, evaluations using PCC or –log10(p value of PCC) (**Fig S2C**) consistently showed that stMLnet exhibited higher LR-target correlations than CytoTalk and random pairing.

Closer cells have a larger probability to communicate with each other, so intercellular signaling should have a stronger impact on intracellular gene expression. To further inspect LR-target correlations of stMLnet, we divided sender-receiver pairs, based on cellular pairwise distances, into a close group (cell pairs with a distance less than the 25th percentile of all of the pairwise distances) and distant group (cell pairs with distance greater than the 75th percentile of all of the pairwise distances). We set malignant cells as receivers and examined whether the close group had a higher correlation (MI or PCC) between LR signaling activity and the target gene expression than the distant group. The results (**Fig S2D**) show that both MI values and PCC values as well as –log10(p value of PCC) of the predicted 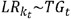 were larger in the close group than in the distant group. These results further verify the reliability of stMLnet in LR-target predictions.

### Analysis of spatially dependent LR signaling in stMLnet

To scrutinize the distance-weighted LR signaling activity, we performed a simulation study based on a mathematical model of ligand diffusion and L-R-TF-Target regulation (see details in **Text S4**) to generate 100 sets of synthetic data (**Fig 3A-B, Fig S3**). We substituted the weighting function of LR signaling quantification in the stMLnet model (i.e., the reciprocal function 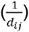; Equation (1)) with the alternative constant (1) or radial basis functions 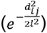. Note that the constant weight corresponds to spatially independent model and the radial basis weight represents the one used by previous methods such as Misty ^25^. We compared the performance of stMLnet with the two counterparts using synthetic data. The simulated spatial expression data were used as input for stMLnet and its variants (without using the prior information of the predefined multilayer network) to infer the regulation of TGs’ expression by LR pairs. The predicted importance scores for LR-TG regulations were benchmarked with the ground truth of the simulated network topology. Various evaluation metrics, including AUROC (area under the ROC curve), AUPRC (area under the Precision/Recall curve), PPV (positive predictive value), Accuracy, Error rate, and MCC (Matthews correlation coefficient), were used for assessment. The evaluation of the 6 metrics on 100 synthetic datasets (**Fig 3C**) showed that the reciprocal function used in stMLnet significantly outperformed the other two variants of weighting functions. These results affirm the rationality and effectiveness of stMLnet in modeling spatial distance-dependent cell communication and gene regulation.

**Fig 3.**
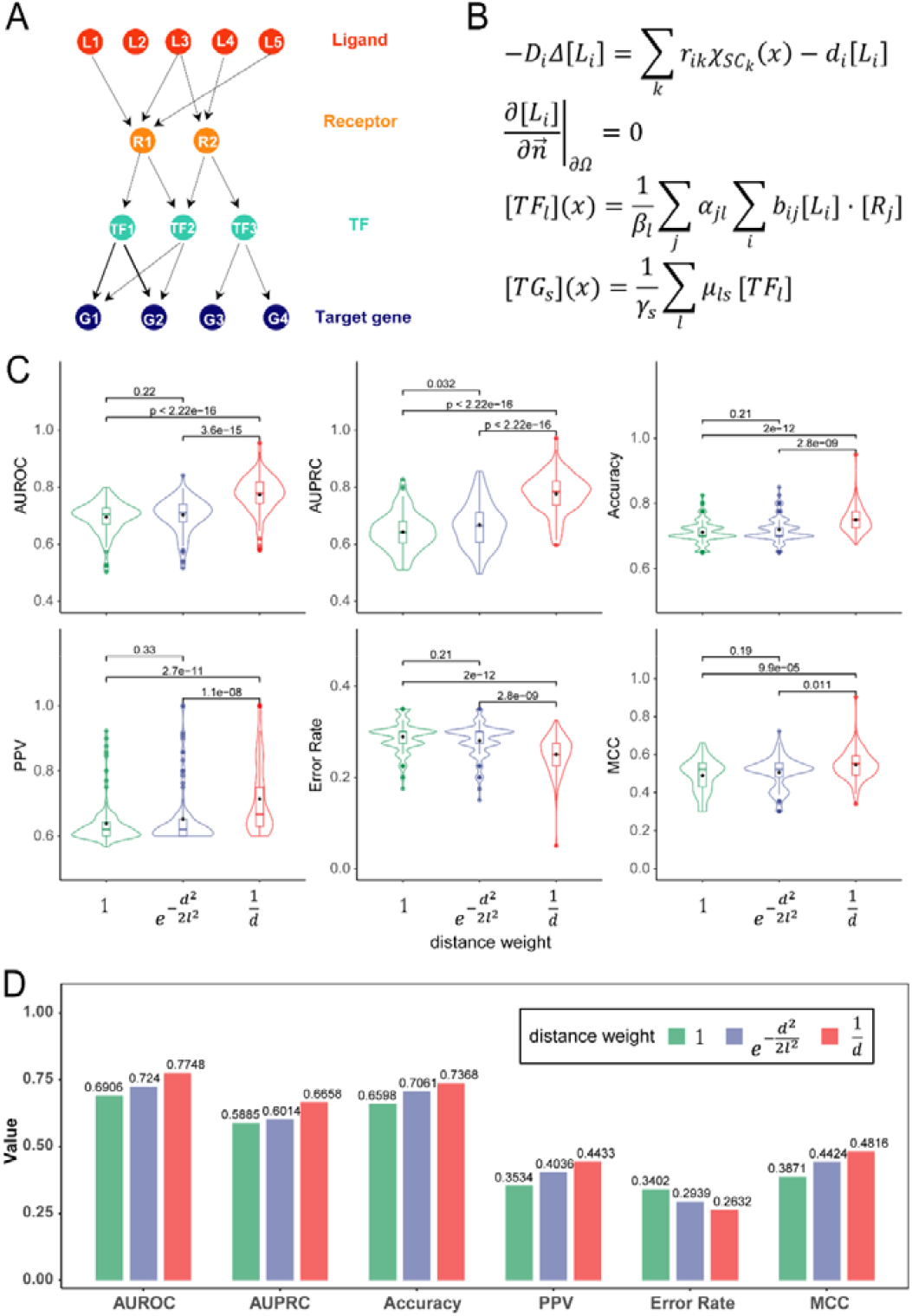
Verifying distance-weighted LR signaling activity in the stMLnet model using synthetic data and perturbation data. The performance of stMLnet in LR-target prediction was compared to that of its variants with modified distance weighting functions in LR activity scoring. The distance weighting functions are the reciprocal function (i.e., 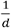) as it is in stMLnet and radial basis function (i.e., 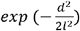) or constant function (i.e., 1) in its two variants, respectively. (**A**) The ground truth of the structure of the simulated multilayer network. (**B**) The mathematical model used for simulation. (**C**) Based on 100 synthetic datasets, the performances of stMLnet with the reciprocal weighting function and the two variants in predicting LR-target regulation were evaluated and compared. The Wilcoxon rank sum test p value was used to assess the statistical significance. (**D**) Based on cell line perturbation data, the performances of stMLnet and the two variants were evaluated and compared. The evaluation metrics (AUCROC, AUCPR, Accuracy, PPV, Error rate, and MCC) were averaged on 7 cell line datasets.

We also compared the prediction accuracies of LR-target regulation based on the above three distance weights using the cell line perturbation data (as used in **Fig 2F**). All of the above evaluation metrics were averaged on the 7 cell line datasets, and stMLnet consistently performed better than the other two variants, favoring the reciprocal distance weight for modeling LR signaling activity (**Fig 3D**).

### Multilayer signaling analysis for CCCs in ST data with high spatial resolution

We next applied stMLnet to several single-cell resolution ST datasets that were generated from different technologies, including seqFISH+, Slide-seq v2, and MERFIS. We first utilized stMLnet to analyze CCCs in the mouse cerebral cortex microenvironment based on a seqFISH+ dataset ^26^ (**Fig 4A**). Abundant and active intercellular communications between different cell types were observed (**Fig 4B**). The LR signaling analysis (**Fig 4C**) showed that SEMA5A-PLXNB3 was the most active upstream LR signaling of OPC (Oligodendrocyte precursor cells), transmitted from L6-eNeuron and L4-eNeuron, respectively (**Fig 4C** upper panel). Furthermore, SEMA5A, LAMC1, and AGRN secreted by OPC, astrocytes, and endothelial cells, respectively, were among the most active ligand signals affecting Olig (oligodendroglia) cells (**Fig 4C** lower panel). Furthermore, multilayer signaling networks (**Fig 4D-E**) depict the regulatory paths from a specific ligand/receptor to downstream TFs and target genes in Oligo or OPC. We found that the LR signaling SEM5A5-PLXNB3 simultaneously existed for mutual communication between Oligo and OPC (**Fig 4E**), consistent with **Fig 4C**. More specifically, SEM5A5 secreted from Oligo cells imposed strong influence on the expression of downstream target genes of OPC (**Fig 4E**), and, in turn, SEMA5A secreted by OPC binds to PLXNB3 on Oligo surface, regulating the expression of multiple target genes (MAG, PCYT2, etc.) through activation of the TF NFKB1 (**Fig 4D**). Collectively, these results suggest a bidirectional feedback loop between OPC and Oligo cells via SEM5A5-PLXNB3-mediated signaling pathways (**Fig 4F**).

**Fig 4.**
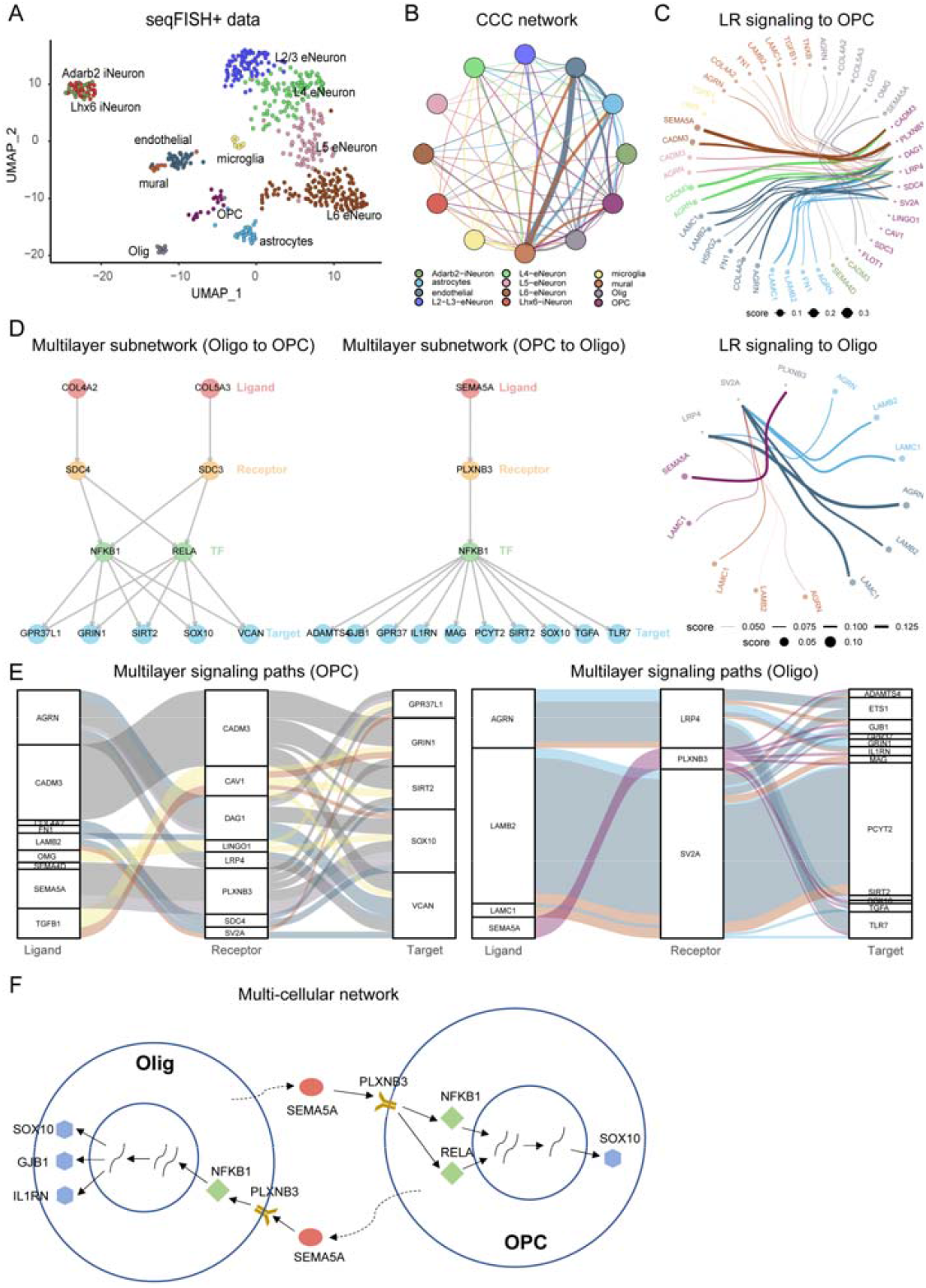
Multilayer signaling analysis for CCCs in seqFISH+ data using stMLnet. (**A**) The cell type annotation in the seqFISH+ dataset of mouse cortex. (**B**) The CCC network. (**C**) The edge bundling plot of LR signaling activity (top ranked). (**D**) The multilayer signaling subnetwork indicates the regulatory paths from a specified ligand/receptor to its downstream TFs and then to target genes. Regulatory paths from Olig to OPC (left panel) and those from OPC to Olig (right panel) are show. Top-ranked target genes according to importance scores are prioritized for visualization. (**E**) The waterfall plot of the multilayer signaling network. Regulatory paths from upstream LR pairs (top ranked) to their downstream targets in Olig (left panel) or OPC (right panel) are shown. The path color indicates the cellular source (sender cells) of the ligand signaling, and the path width represents the importance score of each LR with respect to TG regulation. (**F**) Multiscale CCC network depicting multilayer signaling regulations for a bidirectional feedback loop between OPC and Oligo cells via SEM5A5-PLXNB3 signaling.

We then employed stMLnet to analyze the Slide-Seq V2 data of the mouse hippocampus ^27^ (**Fig 5A**). The CCC network and LR signaling analysis (**Fig 5B-C**) revealed active intercellular signaling communications between the entorhinal cortex and Dentate-Principal-cells and CA1-Principal-cells, which is consistent with the two classical pathways connecting the entorhinal cortex and the hippocampus ^28,29^. Interestingly, the entorhinal cortex may also have active signal transmission with interneurons, as previously reported ^30^. Setting the entorhinal cortex as the sender cells, stMLnet revealed that the upstream LR signaling CALM1/CALM3-KCNQ3 and NCAM1-NCAM1 have the most prominent regulatory potential on the downstream target genes such as CST3, TCF4, ALDOC in CA1-Principal-cells and SST in interneurons, respectively (**Fig 5D**). stMLnet can also be applied to MERFISH data generated by a situ hybridization technology. We analyzed a MERFISH dataset that measured the expression of 161 genes in the mouse brain’s preoptic regions. The dataset consists of 12 spatially adjacent layers along the anterior-posterior axis, and we used four layers of data (layer 3, 6, 9, 12) for CCC analysis. To adapt to the characteristics of low gene coverage of MERFISH technology, we left out Fisher exact test when constructing multilayer signaling network. We found that, despite slight variations in the cell numbers, the CCCs between various cell types remained relatively stable (**Table S3, Fig S4**), implying robustness of stMLnet inference. We visualized the CCC network for layer 9 (**Fig 5E-F**) and found that Excitatory, Ependymal, and other cell types were the primary sender cells, while Endothelial cells, Microglia, and other cell types were the primary receiver cells. Moreover, PNOC-OPRL1 was one of the most active upstream LR signals for dominant cell interactions, such as communications between Astrocyte-Endothelial cells and OP-Mature-Endothelial cells (**Fig 5G**). Furthermore, multilayer network analysis (**Fig 5H-I**) showed that SEMA3C of endothelial cells was regulated by microenvironmental signals from various other cell types including Astrocytes, Excitatory interneuron, and Inhibitory interneuron. SEMA3C has been reported to play important roles in axon growth ^31^. Interestingly, it revealed that both Excitatory interneuron and Inhibitory interneuron can secrete ligand OXT to interact with the vasopressin receptor family (AVPR1A and AVPR2) of Endothelial cells, modulating the activation of the transcription factor ESR1 and regulating the expression of SEMA3C.

**Fig 5.**
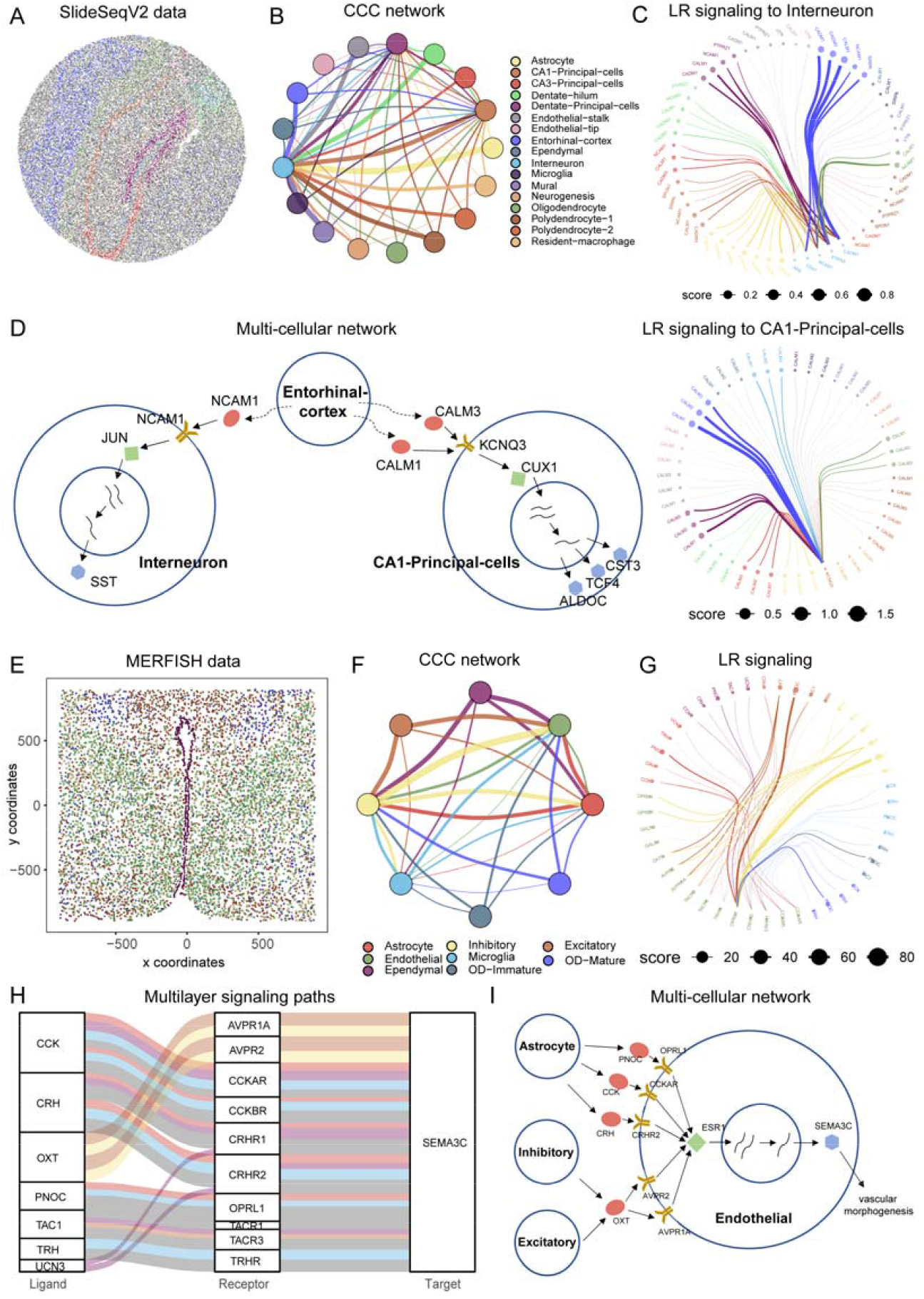
Multilayer signaling analysis for CCCs in Slide-Seq v2 data (A-D) and MERFISH data (E-I) using stMLnet. (**A**) The cell type annotation in the Slide-Seq v2 data of mouse hippocampus. (**B**) The CCC network. (**C**) The edge bundling plot indicates the LR signaling of Interneuron (up) or CA1-Principal-cells (below) transmitted by other types of cells in the microenvironment. (**D**) Multicellular network depicting regulatory mechanism by which Entorhinal-cortex cells regulate Interneuron and CA1-Principal-cells via upstream signals of NCAM1 and KCNQ3, respectively. (**E**) The cell type annotation in the MERFISH data of mouse hypothalamus preoptic region. (**F**) The CCC network. The edge width represents the number of LR pairs. (**G**) The edge bundling plot of LR signaling activity (top ranked). (**H**) The waterfall plot of the multilayer signaling network in Endothelial cells. (**I**) Multiscale CCC network depicting regulatory mechanism by which SEMA5A in Endothelial cells is regulated by upstream signals from other types of cells (e.g., Excitatory, Inhibitory and Astrocytes).

### Application and verification of stMLnet on Visium ST data of gliomas

CSF1R inhibitor (BLZ945) treatment is a macrophage-targeting immunotherapy for gliomas ^32^. However, acquired resistance to CSF1R inhibition often emerges during the treatment, as observed in animal experiments ^33^. To investigate the microenvironment-mediated mechanism underlying resistance to CSF1R inhibition (a macrophage-targeting immunotherapy) in gliomas, we applied stMLnet to a set of Visium ST data ^34^ (**Fig 6A**). The inferred CCC network (**Fig 6B**) demonstrates abundant intercellular LR signaling from glioma cells to macrophages, indicating that glioma cells may substantially impact macrophages’ functions through intercellular interactions. The multilayer networks (**Fig 6C**) further demonstrate the signaling paths downstream of IL34-CSF1R in macrophages (left panel) or those downstream of IGF1-ITGAV in tumor cells (right panel). The transcription factor NFKB1 were found to be involved in the downstream of CSF1R signaling, which is consistent with the experimental studies ^33^. GO/KEGG enrichment analysis for downstream target genes in the multilayer networks (**Fig S5**) revealed several important dysregulated pathways, such as the JAK-STAT pathway, PI3K-AKT pathway and MAPK pathway, induced by glioma-activated CSF1R signaling in macrophages (**Fig S6A, C**) and macrophages-activated ITGAV signaling in tumor cells (**Fig S6B, D**).

**Fig 6.**
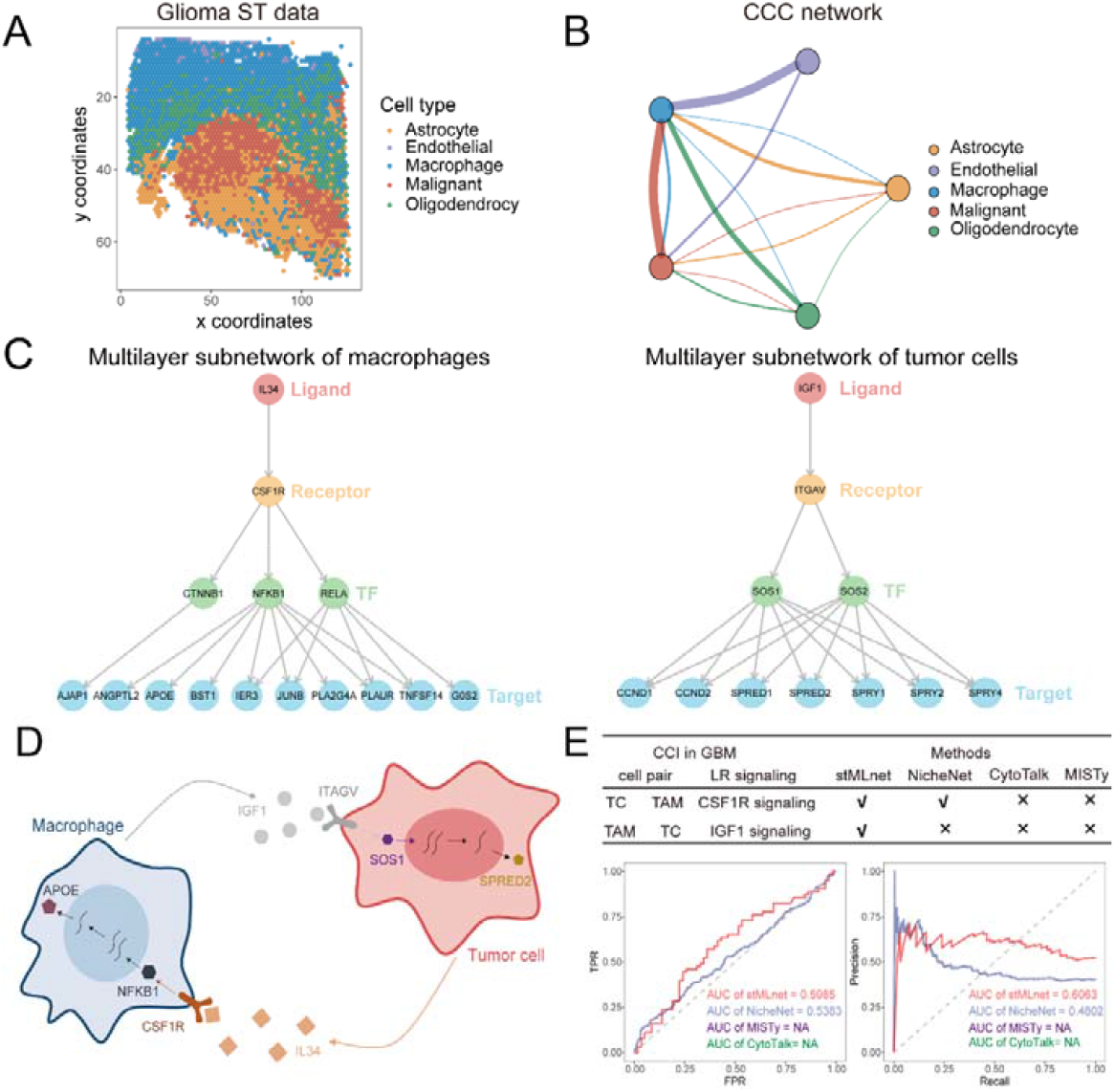
Application and verification of stMLnet on ST data of gliomas. (**A**) The cell types in the ST data. (**B**) The CCC network. (**C**) The multilayer signaling subnetworks downstream of IL34-CSF1R signaling in macrophages (left panel) and that downstream of IGF1-ITGAV signaling in tumor cells (right panel). Only top-ranked target genes are shown for visualization purposes. (**D**) A chart for the inferred interactions between macrophages and glioma cells. (**E**) Verification and comparison of prediction of stMLnet with NicheNet and CytoTalk regarding CSF1R-targets regulation in macrophages using literature evidences (upper panel) or gene expression changes after CSF1R inhibitor treatment in responder mice (lower panel). stMLnet outperformed NicheNet, while CytoTalk and MISTy could not predict out CSF1R signaling (‘NA’).

Among the abovementioned inferred intercellular LRs between glioma cells and macrophages, CSF1R signaling and IGF1 signaling have been validated to be involved in macrophage-glioma interactions and immunotherapy resistance ^33^. stMLnet not only correctly predicted those known ligands or receptors, but also revealed additional signaling pathways (**Fig 6D**), while NicheNet and CytoTalk failed to predict all of the known interactions (**Fig 6E**, upper panel). In addition, to verify the stMLnet prediction regarding CSF1R-targets regulation, a set of RNA-seq data of macrophages isolated from mice (GSE69104) was analyzed. The inferred importance score for CSF1R-targets was compared to the differential expression status (true or false) of the target genes between the CSF1R-responder group (EP) and Veh group. AUCROC and AUCPR were calculated for quantitative evaluation. The CSF1R-targets predictions by NicheNet and CytoTalk were also evaluated for comparison. Eventually, stMLnet achieved higher accuracy than NicheNet, CytoTalk and MISTy (**Fig 6E**, lower panel).

### stMLnet revealed positive cellular feedback circuits and multilayer signaling regulations in a COVID-19 microenvironment

Finally, we applied stMLnet to a set of Visium ST data of COVID-19-infected lung tissue ^35^ (**Fig 7A, Fig S7**) to investigate CCCs underlying the inflammatory response to SARS-CoV-2 infection. The CCC network (**Fig 7B**) showed abundant and active intercellular interactions between various cell types, indicating dysregulated hyperinflammation in the COVID-19-infected lung tissue microenvironment.

**Fig 7.**
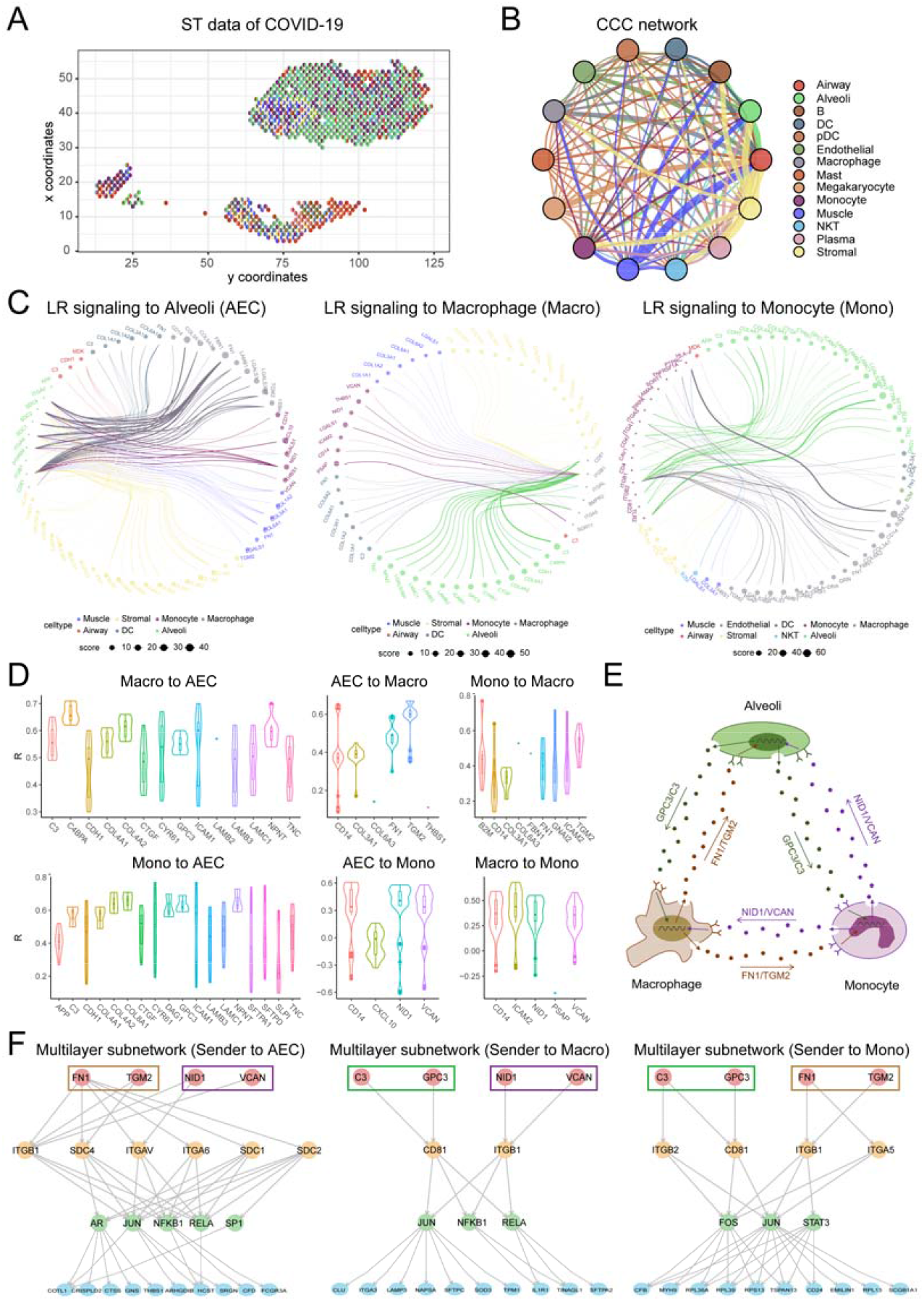
stMLnet revealed positive cellular feedback circuits and multilayer signaling regulations in ST data of COVID-19. (**A**) The ST data of COVID-19 patient. (**B**) The CCC network. (**C**) The intercellular LR signaling network with alveolar epithelial cells (left panel), macrophages (middle panel), or monocytes (right panel) as receiver cells. The node size or edge width indicates the averaged strength of the LR signaling. Several top ranked LR pairs were selected for visualization. (**D**) Correlations between expressions of ligand genes as intracellular targets and their upstream LR signaling activities. Most PCC values (R) were positive except for a small fraction of monocytes. (**E**) Positive feedback circuits between AECs, macrophages, and monocytes. Representative ligands as paracrine cytokines are shown. (**F**) The multilayer signaling paths from the representative paracrine ligands to targets in AECs, macrophages, or monocytes inferred by stMLnet. Top-ranked target genes according to importance scores were prioritized for visualization.

We focused on communications between alveolar epithelial cells (AECs), macrophages, and monocytes, as the latter two are pivotal innate immune cells against SARS-CoV-2 infection ^36^, while AECs express a high level of the SARS-CoV-2 receptor ACE2 ^37^, acting as a master communicator during viral infection ^38^. The LR signaling analysis (**Fig 7C**) depicted paracrine signaling from other cell types to AECs (left panel), macrophages (middle panel), and monocytes (right panel). Interestingly, relatively stronger activities of LR signaling were observed for each pair of the above three cell types. For example, FN1-SDC4 and TGM2-ITGB1 signaling from macrophages to AECs and C3-CD81 and GPC3-CD81 signaling from AECs to macrophages exhibited clearly stronger activities than other LR pairs. Moreover, we found that many ligand genes (e.g., FN1 and TGM2 in macrophages; C3 and GPC3 in AECs; and CD14, NID1, VCAN in monocytes) were also target genes in the multilayer networks of the three cell types (**Table S4**), and that almost all of these ligand genes as targets were positively correlated with their upstream LR signaling (**Fig 7D**). Collectively, these results suggest positive feedback loops between AECs, macrophages, and monocytes through paracrine signaling interactions (**Fig 7E**), which may account for the sustained production and accumulation of inflammatory cytokines and dysregulated hyperinflammation in severe COVID-19 patients ^39^.

Furthermore, the multilayer subnetworks (**Fig 7E, Fig S8**) demonstrate the signaling paths from representative ligands to targets in AECs, macrophages, or monocytes. Several transcription factors, such as NFKB1, RELA, and JUN, were inferred to be involved in the intracellular signaling pathways underlying the above intercellular feedback circuits. These TFs are reportedly critical in regulating the gene expressions of inflammatory cytokines during SARS-CoV-2 infection, and they are targets of FDA-approved drugs ^40^, indicating their therapeutical values for COVID-19 by disrupting the above feedback circuits. In addition, we investigated the biological functions of the above intercellular feedback circuits by performing functional enrichment analysis for their intracellular target genes. The GO and KEGG pathway enrichment (**Fig S9**) demonstrated that biological processes or pathways related to COVID-19, immune response, cell adhesion, and the extracellular matrix (ECM) were significantly enriched for communications among AECs, macrophages, and monocytes. These results indicate the important roles of the abovementioned intercellular feedback loops in pulmonary injury and immune disorders in response to SARS-CoV-2 infection, and they are consistent with the results of previous studies ^41,42^.

## Discussion

We have described stMLnet, a tool to dissect multiscale CCCs in ST data. It combines knowledge graph, statistical inference, mechanistic modeling, and machine learning to construct a multilayer signaling network, infer LR signaling activity, and predict LR-target genes regulation. stMLnet leverages spatial information in the ST data to quantify intercellular signaling activity and, more importantly, connect extracellular signals to intracellular gene expression. stMLnet can be used to depict the microenvironmental regulation of gene expression by prioritizing upstream regulators for a given set of target genes or inferring downstream pathways and regulatory networks for a given ligand or receptor.

One of the challenges in CCC inference is the lack of appropriate benchmarking, which impedes the methodological development and practical applications of such tools. As such, the predictions derived from different methods or tools sometimes differ from each other, and their prediction accuracies have not been well tested. In this study, we collected cell type-specific perturbation datasets for quantitative benchmarking LR-targets predictions. Although *in vitro* data are not the perfect gold standard, to the best of our knowledge, cell line perturbation data have been the best choice for benchmarking ligand-target predictions for now, as differential responses of target genes to ligand or receptor perturbations characterize the potential regulations between them. In addition, we utilized *in vivo* perturbation data of cells from glioma-bearing mice to verify CSF1R downstream signaling. Among the existing methods ^12-15,18,19,25,43-54^ (**Table S1**), we chose NicheNet, CytoTalk, and MISTy as competitors for benchmarking stMLnet, as they can output prediction scores of ligand-target regulations that can be compared to the ground truth (differential expression of targets in response to ligand/receptor perturbations). The results showed that stMLnet outperformed the other methods on multiple datasets.

The advances of stMLnet are as follows. First, the L-R-TF-TG multilayer network modeling framework is adequate and effective in depicting intercellular communication-mediated gene expression. The multilayer network method evaluates the upstream activities using the downstream responses, which may reduce false-positive predictions. To fulfill this objective, we integrate carefully curated prior network database information with context- and cell type-specific expression data to discover potential causal relationships between the upstream signaling and the downstream targets by employing the random walk algorithm and statistical inference method (e.g., Fisher’s exact test). Second, the spatial information in the ST data is leveraged to mechanistically quantify LR signaling activities based on a mathematical diffusion model of microenvironmental ligands. This leads to a parameter-free reciprocal relationship between effective ligand signals receipted by the receiver cells and cell–cell physical distance, which, although simplified, avoids unfeasible estimation or calibration of ligand-specific parameters (e.g., diffusion rate and degradation rate). Third, we employ an explainable random forest regression model to measure the contribution of each ligand or receptor or LR pair in regulating target gene expression.

Admittedly, our study had some limitations. For example, our method makes several simplified assumptions. In our diffusion model, we approximate the ligand concentration around the sender cells with ligand gene expression by assuming a positive correlation between them. Similarly, regarding the LR signaling activity, we use receptor gene expression to substitute its protein level in the receiver cells. Additionally, the TF-target information comes from prior databases, which are not cell-type-specific. These drawbacks can be addressed by integrating ST data with emerging single-cell multi-omics data, such as single-cell proteomics data and single-cell ATAC sequencing data, thanks to the fast development of sequencing technology. In future studies, we will develop novel multiscale models that integrate multi-omics data across multiple layers to make better inferences of CCC and gene–gene regulations.

## Conclusions

In summary, stMLnet, a method for modeling spatially dependent intercellular signaling and intracellular regulations, serves as a valuable tool for analyzing multiscale signaling and functioning of CCCs in spatial transcriptomics data.

## Methods

The proposed method stMLnet mainly encapsulates four components (**Fig 1**), i.e., constructing prior network databases based on multiple data sources, inferring multilayer signaling networks, calculating distance-dependent LR signaling activity, and quantifying regulatory relationships between upstream L/R and downstream target genes in the inferred multilayer network.

### Collection and integration of prior network information

To infer inter- and intra-cellular signaling networks, we collected multiple data sources of molecular interactions as prior network information (**Text S1**). We processed and integrated them into prior knowledge databases of stMLnet at three scales of signaling transduction: LigRecDB (ligand-receptor prior database), TFTGDB (TF-target gene prior database), and RecTFDB (receptor-TF prior database) (**Fig S10**) (**Table S5**).

#### LigRecDB

The ligand-receptor interaction information was collected from connectomeDB2020 ADDIN^55^, iTALK ^15^, NicheNet ^18^ and CellChat ^14^. Most of the ligand-receptor pairs deposited in these databases are evidenced by the literature reports or supported by multiple other databases. Ultimately, we curated a ligand-receptor database LigRecDB that consists of 3659 pairs of non-redundant LR interactions, including 920 ligands and 751 receptors.

#### TFTGDB

The interaction information between TF and target genes was collected from TRRUST ^56^, HTRIdb ^57^, RegNetwork ^58^ and GTRD ^59^. TRRUST and HTRIdb deposit many literature-supported TF-target genes interactions. RegNetwork integrates 25 previously existing databases including predicted/experimental transcriptional regulatory interactions. Ultimately, we collected 373501 pairs of TF-target gene interactions, with 525 TFs and 23021 targets, which compose the TFTGDB database.

#### RecTFDB

To infer links from receptors to TFs, we used R package graphite ^60^ to extract information of intracellular signaling pathways or molecular interactions from 8 existing databases. Based on such information, we constructed directed weighted graph and employed a random walk with restart algorithm ^61^ to infer receptor-TF links (**Text S2**) and the hyper-parameters were optimized using perturbation-expression data of 69 cell lines involving 27 ligands and 13 receptors (**Fig S11**). At last, 17450 non-redundant receptor-TF links were obtained, comprising 751 receptors and 525 TFs, which compose the RecTFDB database.

The comparison of the prior databases of stMLnet with those of NicheNet ^18^ and Omnipath ^62^ is described in **Text S5**.

### Inference of multilayer intercellular and intracellular signaling networks

The multilayer signaling network comprises of four layers of signaling molecules (i.e., ligand, receptor, TF and target genes) and three coherent sub-networks (i.e., LR signaling, receptor-TF pathways, and TF-target gene interactions). Below we describe details of the inference of structure of the multilayer networks.

#### Constructing ligand-receptor subnetwork

Based on the LigRecDB, we selected highly expressed ligands and receptors as potential L-R signaling pairs. Differentially-expressed genes (DEGs) that are specifically expressed in the sender cells (by Seurat ‘FindMarkers’, default parameters: min.pct = 0.1, logfc = 0.25) constituted the set of highly-expressed ligands (LIGs). The highly-expressed genes in the receiver cells with average expression level greater than a threshold (e.g., 0.05 as default) and expression percentage more than a threshold (e.g., 10% cells as default) constituted the set of highly-expressed receptors (RECs). The resulting ligand-receptor subnetwork was denoted as N_LIG-REC_.

The above parameters for selecting LIGs and RECs could be adjusted by the users for the specific problems. For the ST datasets of breast cancer and COVID-19, we used the default parameters as described above; for the scRNA-seq dataset of glioma, the RECs were analyzed using the default parameters while the LIGs were selected using another set of parameters: min.pct = 0.05, logfc = 0.1, considering the sparsity of the scRNA-seq data.

#### Constructing TF-target gene subnetwork

The users can input a set of genes of interest as potential target genes, which can also be set as differentially expressed genes (DEGs) (via, e.g., ‘FindMarkers’ function in Seurat) or interaction-changed genes (ICGs) (via, e.g., ‘findICG’ function in Giotto, default). The set of input target genes was denoted as aTGs, other genes were denoted as naTGs. We used Fisher’s exact test to calculate a p-value for each *TF_i_* as follows,

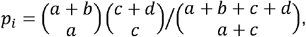

where 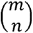 represents the binomial coefficient. We denoted TGs to represent the gene regulated by *TF_i_* and nTGs the genes that are not regulated by *TF_i_*. *a* |*aTGs* ⋂ *TGs* | represents the number of genes in the intersection between input target genes and TF_i_-regulated target genes. *b* = | *aTGs*| − *a* represents the number of genes that are in the input target gene set but are not regulated by *TF*_*i*_. *c* | *TGs* | − *a* represents the number of genes that are regulated by *TF_i_* but are not belong to the input target gene set. *d =* | *naTGs* ⋂ *nTGs*| represents the number of genes that are neither in the input target gene set nor regulated by *TF_i_*. If the Fisher’s exact p-value *p_i_* ≤ 0.05, then the *TF_i_* is considered to be significantly activated. All the activated TFs constitute a set aTFs. As a result, we get the TF-target gene subnetwork *N*_*TF-TG*_ by paring TFs in aTFs with genes in aTGs.

#### Constructing receptor-TF subnetwork

The construction of receptor-TF subnetwork is similar to that of TF-target gene subnetwork. We employed Fisher’s exact test to calculate a p-value for each receptor *REC*_*i*_ as follows,

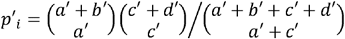

We denoted *TF_s_* to represent the TFs regulated by *REC_i_* and *nTF_s_* the genes that are not regulated by *REC*_*i*_. *a* ′ = | *aTFs* ⋂ *TFs*| represents the number of TFs in the intersection set between the activated TFs and *REC_i_*-regulated TFs. *b*′ = |*aTFs*| − *a* ′ represents the number of TFs that are activated but are not regulated by *REC_i_*. *c*′ = |*TFs*| − *a* ′ represents the number of TFs that are regulated by *TF_i_* but are not activated. *d =* |*naTFs* ⋂ *nTFs*| represents the number of TFs that are neither activated nor regulated by *REC_i_*. If the Fisher’s exact p-value *p*′ _*i*_ ≤0.05, then the *REC_i_* is considered to be significantly activated. All the activated receptors constitute a set aRECs. As a result, we get the receptor-TF subnetwork *N_REC-TF_* by paring receptors in aRECs with TFs in *aTF* based on the RecTFDB database.

#### Integrating subnetworks into a multilayer network

The above subnetworks were connected together into a multilayer signaling network *N_LIG−REC−TF−TG_* by pruning the nodes and links that are not the downstream of the activated LR signaling or activated TFs.

### Modeling spatially dependent ligand-receptor signaling activity

To quantify signaling activities of LR pairs, we employed a mechanistic modeling approach by considering ligand diffusion in the microenvironment. Assume that the spatial coordinates of cell *i* and cell *j* are (*x*_*i*,_ *y*_*i*,_ *z*_*i*_) and (*x*_*j*,_ *y*_*j*,_ *z*_*j*_), respectively. The Euclid distance between cell *i* and cell *j* is 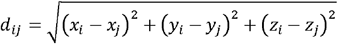. Denote *LR^k^* as the *k*-th pair of ligand-receptor, and 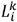 and 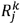 the corresponding ligand expression in the *i*-th cell and receptor expression in the *j*-th cell, respectively.

The ligand is a type of cytokine, which can diffuse in the microenvironment following release by the sender cells. The spatial-temporal distribution of the ligand concentration *u (x,y,z)* during diffusion can be described by a partial differentia equation (PDE) as follows,

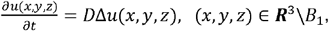

where Δ is the Laplace operator, defined as 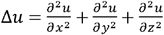. *D* is the diffusion coefficient. *B*_1_ represents unit ball indicating the sender cell. We assume that the diffusion of the ligand is relatively fast and the above equation quickly reaches to the steady-state. Therefore,

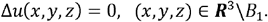

We further assume that the diffusion of the ligand across the local microenvironment is homogenous, and thereby the solution to the above equation is radially symmetric, i.e., *u(x,y,z)* = *u(r)*, where, 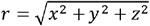. Consequently, the above steady-state diffusion model could be simplified to an ordinary differential equation (ODE):

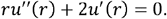

Solving the above ODE, we get

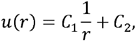

where *C*_1_ and *C*_2_ are constants. Assume that the ligand concentration is *u*_1_ along the boundary of *B*_1_ and 0 far away. So we impose the boundary conditions *u*(1) = *u*_1_, *u* (∞) = 0. As such 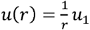. We used the ligand expression in the sender cell 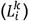 as a proximity of *u*_*1*_. Therefore, the signaling strength of 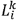 received by the *j*-th cell is 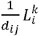.

Furthermore, based on the law of mass-action, the signaling strength of the *k*-th LR pair activated at the *j*-th cell could be defined as

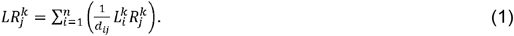

These equations quantitatively describe the influence of cell distance on ligand-receptor signaling intensity, so that the ligands from the sender cells with different distances react with the receptor of the receiver cell in different extent. As a result, the samples of LR signals (i.e., receiver cells) are consistent with those of the corresponding target genes, allowing mapping LR signal to the intracellular target gene expression.

### Random forest regression for LR-target regulation

Assume *m* target genes (i.e., *G*_*1*_, *G*_*2*_, …, *G*_*m*_) in the inferred multilayer network and *n*_t_ pairs of LRs linked to each target gene *G*_t_ (i.e., 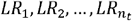). To decipher quantitative regulation relationship between LR pairs and the target genes, we divided this task into *m* sub-problems by decoding the following functions:

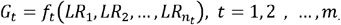

Due to complex regulation relationship between LRs and the target gene, the above *f_t_* is generally non-linear. We thus employed explainable tree-based regression to learn *f_t_*. We constructed random forest regression model for each of the *m* target genes (**Fig 1E**). We used the signaling activities 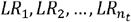 (calculated from Equation (1)) across the receiver cells as input to predict the expression of the target gene *G_t_*.

### Importance ranking of the upstream regulators

Based on the random forest model, we could calculate the importance score of the *k_t_*-th feature 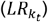 contributing to the expression of target gene *G_t_*, and thereby rank LR pairs. Moreover, we also refined the permutation importance score to account for the disassembled importance of L or R alone, which was referred to partial importance score.

#### Importance score of LR pairs

We calculated permutation-based feature importance for each LR pair by evaluating mean-squared-error (MSE) between the random forest (RF) model prediction and the true values on out-of-bag (OOB) samples and comparing the change of the MSE before and after the permutation of the LR activities. If the MSE is largely increased after LR permutation, then the LR pair is viewed to be important with respect to regulating the target gene expression. To integrate the computed importance scores for the *m* target genes and avoid the bias originated from the variance in gene expression ^63^, we normalized the input and the output (z-score) of the random forest regression model before running it. Consequently, the importance scores computed from the *G_t_* model were denoted as

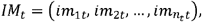

where 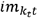 represents the importance score of the *k_t_*-th LR pair for the *G_t_* expression. Integrating the importance scores computed from the *m* models, we get

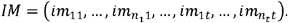

As such, we could compare the importance scores of LR pairs across different regression models.

#### Partial importance score of L or R

The above importance score was defined for the combination of the ligand and the receptor. To decompose the importance score for the ligand or receptor alone in each LR pair, we modified the above permutation-based feature importance method to account for the partial importance score (PIM) of the upstream regulator molecules (ligand or receptor, denoted as *v_q_, q* = 1,2,…, *q_t_*) for the target gene *G_t_*. Specifically, the PIM was computed as follows.

1. For each regression tree, compute the MSE using the OOB data, denoted as *errOOB*_1_;
2. Randomly permute the values of the features (LR pairs) involving *V_q_* on all the OOB samples and then compute the MSE using the OOB data, denoted as *errOOB*_2_;
3. For the random forest with *N* trees, the PIM of the upstream regulator molecule *v_q_* could be calculated as

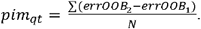

Ultimately, the partial importance scores of the upstream regulators computed from the random forest model *f_t_* were

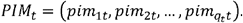

According to the ranking in the above *PIM_t_*, we could compare the regulatory capabilities of different upstream regulatory molecules for the target gene *G*_*t*_. On the other hand, integrating the partial importance scores computed from the *m* models, we get

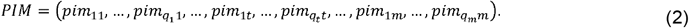

This enables us to compare the regulatory capabilities of a given upstream regulators with respect to different target genes.

### ST datasets

The following ST datasets were used in this study for analysis: seqFISH+ data of mouse cortex ^26^, Slide-seq v2 data of mouse hippocampus ^27^, MERFISH data of mouse hypothalamus preoptic region ^64^, and Visium data of breast cancer ^24^, glioma ^34^ and COVID-19 ^35^. The collection and processing of these datasets are described in **Text S6**. The detailed information of cell line perturbation datasets is provided in **Table S6**.

## Supplementary information

Supplementary information is available online.

**Text S1**. Collection and integration of prior network information.

**Text S2**. Inference and optimization of Receptor-TF regulatory matrix.

**Text S3**. Implementation of CytoTalk, NicheNet and MISTy.

**Text S4**. Simulation study.

**Text S5**. Comparison of prior databases.

**Text S6**. Dataset collection and processing.

**Figure S1**. Enrichment analysis for the multilayer signaling networks of breast cancer.

**Figure S2**. Verifying stMLnet based on LR-target correlations.

**Figure S3**. Illustration for the simulation study.

**Figure S4**. Comparison of cell communication in different layers of MERFISH data

**Figure S5**. The waterfall plot of the multilayer signaling network for the cellular feedback circuits inferred from the Glioma ST dataset.

**Figure S6**. Enrichment analysis for the multilayer signaling networks of glioma.

**Figure S7**. The cell type annotation for the scRNA-seq data of COVID-19.

**Figure S8**. The waterfall plot of the multilayer signaling network for the cellular feedback circuits inferred from the COVID-19 ST dataset.

**Figure S9**. Heatmap of functional enrichment for cellular feedback circuits inferred from the COVID-19 ST dataset.

**Figure S10**. The collection and sorting of prior knowledge database.

**Figure S11**. Comparison and parameterization of prior knowledge database.

**Table S1**. Summary and comparison of cell communication inference methods.

**Table S2**. The perturbation cell line datasets of breast cancer.

**Table S3**. Comparison of potential signals in different layers of MERFISH data

**Table S4**. Correlations between ligand genes and their upstream LR regulators involved in cellular feedback circuits in COVID-19.

**Table S5**. Statistics of information in the constructed prior databases LigRecDB, RecTFDB and TFTGD5B.

**Table S6**. The detailed information of cell line datasets.

## Data availability

The MERFISH data and the seqFISH+ data were download from the spatial-datasets project of giotto, via [https://github.com/drieslab/spatial-datasets/tree/master/data/2018_merFISH_science_hypo_preoptic] and [https://github.com/drieslab/spatial-datasets/tree/master/data/2019_seqfish_plus_SScortex], respectively. The Slide-seq v2 data was download from SeuratData package [https://www.biorxiv.org/content/10.1101/2020.03.12.989806v1]. The ST data of breast cancer was downloaded from the 10X Genomics website (https://support.10xgenomics.com/spatial-gene-expression/datasets/1.0.0/V1_Breast_Cancer_Block_A_Section_1). The ST data of glioma was downloaded from https://github.com/theMILOlab/SPATAData. The ST data of COVID-19-infected lung tissue was downloaded from Mendeley (https://doi.org/10.17632/xjtv62ncwr.1). The scRNA-seq data of breast cancer, the scRNA-seq data of glioma, and the bulk RNA-seq data of macrophages were downloaded from the NCBI GEO database (GSE118389, GSE84465, and GSE69104, respectively). The scRNA-seq data of the COVID-19 was downloaded from Synapse (https://www.synapse.org/#!Synapse:syn21041850). The cell line gene expression datasets were downloaded from the NCBI GEO database, with the access numbers listed in **Table S6**. The simulation data is available from a public repository of Github (https://github.com/SunXQlab/stMLnet-simulation).

## Code availability

The source code for the data analysis in the manuscript is available from Github (https://github.com/SunXQlab/stMLnet-AnalysisCode). The R package of stMLnet is publicly available from Github (https://github.com/SunXQlab/stMLnet).

## Authors’ contributions

J.C. performed research, analyzed data, and wrote the draft; L.Y. analyzed data and assisted the software development; Q.N. participated in discussion and edited the paper; X.S. designed study, performed research, analyzed data, and wrote the paper.

## Acknowledgements

We would like to acknowledge members in the Sun Lab at SYSU for valuable discussion and technical assistance.

