## Supplementary material for "Modeling and inference of spatial intercellular communications and multilayer signaling regulations using stMLnet": Text S1-S6; Figure S1-S11; Table S1-S6

**Outline**

**Text S1**. Collection and integration of prior network information.

**Text S2**. Inference and optimization of Receptor-TF regulatory matrix.

**Text S3**. Implementation of CytoTalk, NicheNet and MISTy.

**Text S4**. Simulation study.

**Text S5**. Comparison of prior databases.

**Text S6**. Dataset collection and processing.

**Figure S1.** Enrichment analysis for the multilayer signaling networks of breast cancer.

**Figure S2**. Verifying stMLnet based on LR-target correlations.

**Figure S3**. Illustration for the simulation study.

**Figure S4**. Comparison of cell communication in different layers of MERFISH data

**Figure S5.** The waterfall plot of the multilayer signaling networks in the Glioma ST dataset.

**Figure S6**. Enrichment analysis for the multilayer signaling networks of glioma.

**Figure S7**. The cell type annotation for the scRNA-seq data of COVID-19.

**Figure S8**. The waterfall plot of the multilayer signaling network for the cellular feedback circuits inferred from the COVID-19 ST dataset.

**Figure S9**. Heatmap of functional enrichment for cellular feedback circuits inferred from the COVID-19 ST dataset.

**Figure S10**. The collection and sorting of prior knowledge database.

**Figure S11**. Comparison and parameterization of prior knowledge database.

**Table S1**. Summary and comparison of cell communication inference methods.

**Table S2**. The perturbation cell line datasets of breast cancer.

**Table S3**. Comparison of potential signals in different layers of MERFISH data.

**Table S4**. Correlations between ligand genes and their upstream LR regulators involved in cellular feedback circuits in COVID-19.

**Table S5.** Statistics of information in the constructed prior databases LigRecDB, RecTFDB and TFTGD5B.

**Table S6.** The detailed information of cell line datasets.

**Supplementary References**

**Text S1. Collection and integration of prior network information**

***LigRecDB***

The LigRecDB database in stMLnet collected the information from various databases including CellChat, connectomeDB, NicheNet and iTALK. These four databases deposited information of interactions between ligands and receptors, which covered the majority of data sources such as protein-protein interaction database, pathway database, gene regulatory database and literature-mining.

CellChat [1] collected four categories of ligand-receptor interaction information (interactions, complexes, co-effectors, and annotations) based on KEGG and literature-mining. After converting the interaction involving complexes into the interaction between molecules by using the information from complexes, we obtained 1921 molecular interactions from CellChat, including 514 ligands and 439 receptors.

Based on Ramilowski et al. [2], ConnectomeDB2020 [3] integrated some databases that have been embedded in several cell-cell communication tools, including CellphoneDB [4], RNA-Magnet [5], SingleCellSignalR [6], ICELLNET [7], etc. These ligand-receptor interactions are all supported by literatures. We extracted 2293 pairs of ligand-receptor interactions from ConnectomeDB2020, involving 829 ligands and 690 receptors.

iTALK [8] was derived from text mining and divided ligand-receptor interactions into "checkpoint interaction", "cytokine interaction", "growth factor interaction" and "other interaction". To minimize information redundancy, we excluded interactions coming from connectomeDB2020 and Ramilowski et al. database when dealing with iTALK. As a result, 2440 ligand-receptor interactions were obtained from iTALK, involving 692 ligands and 664 receptors.

NicheNet [9] collected ligand-receptor interaction information from 14 data sources, including text mining, pathway databases, drug databases, and molecular interaction databases. We excluded some relatively unreliable ligand-receptor interactions when dealing with NicheNet, including interactions coming from Ramilowski et al. database but filtered in connectomeDB2020 and interactions predicted by NicheNet through molecular interaction database and GO database. We screened out 2010 ligand-receptor interactions involving 536 ligands and 509 receptors from NicheNet.

We integrated the ligand-receptor interactions extracted from the above fours databases and only retained the interactions with HGNC gene correspondence. Finally, we collected 3860 non-redundant ligand-receptor interactions, containing 972 ligands and 863 receptors. Some details of interactions were also recorded, for instance, the primary sources, the secondary sources, supporting literatures and occurrence times (**Fig S10A-B** and **Table S1**).

***TFTGDB***

The TFTGDB database in stMLnet collected interactions between transcriptional factors (TFs) and target genes (TGs) from TRRUST, RegNetwork, HTRIdb and GTRD.

TRRUST collected 9396 pairs of TF-TG interactions through literature mining, and labeled specific regulatory relationships as ’activated’, ‘inhibited’ and ‘unknown’. We filtered miRNA interactions and contradictory regulator interactions, ultimately preserving 8427 regulatory interactions.

RegNetwork was developed based on 25 databases, 17 of which provide regulatory relationship information and 8 of which are auxiliary databases. We downloaded regulatory interaction information in RegNetwork website, and filtered miRNA-mediated interactions and repeated interactions, and finally retained 155426 pairs of regulatory interactions.

HTRIdb collected predicted and experimentally validated TF-TG interactions from classic regulatory databases such as JASPAR, TRANSFAC, TRED, and TRRD, as well as literatures. With the same filtering criteria (excluding miRNA-mediated interactions and repeated interactions), we obtained 18160 transcriptional regulatory interactions between 283 TFs and 11,886 TGs.

GTRD interactive website was used to obtain regulatory binding sites near the transcription promoter. With the same filtering criteria (excluding miRNA-mediated interactions and repeated interactions), we obtained 588238 pairs of TF-TG interactions between 569 TFs and 6319 TGs.

Integrating the transcriptional regulation information extracted from the above four databases, we collected 754270 non-redundant TF-TG interactions, involving 1712 transcription factors and 23,836 target gens. We also recorded the following information for each interaction: the primary data sources, secondary data sources, literature support information and occurrence times (**Fig S10A-B** and **Table S1**).

***RecTFDB***

Receptor-transcription factor interactions are mainly collected from eight pathway-related databases by using R package graphite. The component members and topological structure of each pathway were extracted to form three network modes including protein-protein interaction, metabolite-metabolite interaction, and protein-metabolite interaction. Based on the following rules, we obtained the interactions between protein molecules in the pathway using ‘edges’ function: (1) for complex complexes consisting of multiple protein molecules, it was assumed that there are interactions between each protein molecules; (2) For the metabolite-mediated interaction between two proteins (protein A→metabolite C→protein B), we reserved only interaction between proteins (protein A→protein B); (3) For the molecular interaction (A→B) of “Process (Binding /association)” or “undirected” type, we added the reverse interaction direction B→A. A total of 398,809 non-redundant protein interactions were extracted, consisting of 11,354 proteins ranging from upstream regulatory molecules (ligands and receptors) to intermediate molecules (kinases, intermediates, etc.) and downstream molecules (transcription factors and target genes) (**Fig S10A-B** and **Table S1**). Among these proteins, there were 762 receptors on 1145 transcription factors.

Unlike ligand-receptor interactions or transcription factor- target interactions, the regulations between receptors and transcription factors are not directly interacted but indirectly transmitted through the assistance of multiple signal molecules. Moreover, the interaction information collected from the databases is usually uncomprehensive and some of crosstalked pathways are often separately deposited in different databases. For example, receptor *a* in pathway *A* regulates intermediate signal molecule *b* (database 1), while intermediate signal molecule *b* in pathway *B* influences the activation of downstream transcription factor *c* (database 2). In this situation, by observing pathway *A* and pathway *B* separately, the regulatory potential between receptor *a* and downstream transcription factor *c* will be neglected. To resolve the above issues, we employed the pathway information of the above 8 databases to construct an integrated weighted directed graph to evaluate the regulatory potential between receptors and transcription factors. The direction was determined according to the direction of signal transmission, and the weight was determined according to the occurrence times of interactions in the databases. We assumed that receptor-transcription factor pairs with higher regulatory potential are closer to each other in the weighted directed graph than that with lower regulatory potential, and that the closer the distance is, the greater the potential of regulation (the distance is affected by the network topology and node attributes). In most cases, there is no direct connection between the receptor node and the transcription factor node. We can use the possibility of connection between two nodes to express the regulatory potential between the receptor and the transcription factor, which is analogy with the link prediction problem, so we use random walk with restart algorithm to predict receptor-transcription factor linking.

The sketch of the approach to constructing our RecTFDB database is as follows. We first constructed a weighted directed signal network with protein as node and interaction as edge. Each receptor was used as the starting point (seed node) for randomness. The receptor might return to itself with probability *r* (restart probability), or randomly wandered to nearby reachable nodes with 1-*r* probability. After multiple walks or resets, the probability of the receptor reaching to any other node in the graph would stabilize. As such, we obtained a probability matrix of 762 receptors regulating 1145 transcription factors, with each value in the matrix representing the potential of a pair of receptor-transcription factor interaction. These interactions were mapped to HGNC genes and only annotated ones were retained. Finally, a threshold of regulatory potential was determined to further filter some interactions based on an optimization model by fitting it to the collected cell line perturbation data. Consequently, RecTFDB was constructed, consisting of 17,450 non-redundant receptor-transcription factor interactions (751 receptors and 525 transcription factors). LigRecDB and TFTGDB were updated accordingly based on layer-shared receptors and transcription factors, we finally obtained 3659 non-redundant ligand-receptor interactions (920 ligands and 751 receptors) and 373501 non-redundant transcription factor-target gene interactions (525 transcription factors and 23021 target genes), respectively.

The details of receptor-TF link inference are described below (**Text S2**).

**Text S2. Inference and optimization of Receptor-TF regulatory matrix.**

***Weighted directed graph-based random walk algorithm***

To predict receptor-transcription factor pairing, we constructed a directed weighted graph $G\left( V, E_{V} \right)$ based on the available data sources of signaling pathways. $V=\left\{ v_{1},\cdots,v_{n} \right\}$ is the set of the interacting molecules, and $E_{V}$ is the set of the observed interactions between the upstream molecule $v_{i}$ and the downstream molecule $v_{j}$. An adjacent matrix $W$ was also constructed with weight $w_{ij}$ defined as the occurrence number of the interaction ($v_{i}$→$v_{j}$) recorded in the databases.

The link probability between receptor and transcription factor can be considered as a link prediction problem in a graph. We used the random walk with restart (RWR) algorithm [10, 11] to calculate the probability of link generation between each pair of nodes that are not directly connected. The formula for the RWR is as follows:

$p_{t+1}^{T}=\left( 1-r \right)Mp_{t}^{T}+rp_{0}^{T}$,

where $p_{0}$ is initial probability vector for nodes. Given a set of source/seed nodes $S\subset V$, the value of $p_{0}$ is 1 for the seed nodes (i.e., $\frac{1}{\left| S \right|}$) and 0 for other nodes, respectively. $p_{t}$ and $p_{t+1}$ are probability vectors of nodes at time *t* and *t*+1, respectively. The *i*-th element of $p_{t}$ represents the probability of the walker being at node $v_{i}$. $M$ represents transition probability matrix, describing the random walk probability for each pair of nodes, defined as $M_{ij}=\frac{w_{ij}}{\sum_{j} w_{ij}}$. $r\in(0,1)$ is a restart-probability representing the probability of a walker going back to the source nodes. When the above iteration converges, $p_{t}$ could approximately represent the closeness or importance score with respect to the node of interest.

The main idea of RWR algorithm is that a random particle that starts from the seed node *x* can transmits to its neighbor with probability 1-*r* or return to the seed node *x* with probability *r*. After several iterations, the probability of the particle staying at any other node tends to be stable and is related to the seed node initially selected. We assume that the seed node is the receptor, and when it reaches the stable state after RWR process, we can get the probability of the receptor linking with any other node (some are TF nodes). This probability was used to represent the potential regulatory ability of the receptor on the downstream gene in the protein interaction graph constructed based on prior knowledge.

***Determination of restart-probability r***

For the above graph $G(V, E_{V})$, $E_{U}$ represents the set of all possible links between each pair of nodes in $V$. $E_{N}$ denotes a set of unobserved links that exist in $E_{U}$ but not in $E_{V}$. $E_{M}$ stands for missing links (such as receptor-transcription factor pairing) that cannot be detected due to incomplete collection or limitation of current technology. The task of link prediction is to find out the missing links for short.

We used 3-fold cross-validation and random sub-sampling validation [12] to determine the restart-probability *r* and to test the algorithm’s accuracy. The set of observed links $E_{V}$ was divided into three subsets. Each time two subsets were used as training set $E_{T}$, and the remaining one was used as validation set $E_{P}$. Then we repeated the process 3 times, with each of the 3 subsets used exactly once as the validation set. We performed the RWR process on the graph $G'$ constructed from the training set $E_{T}$ and the RWR index would give a score of missing links ${E'}_{M}$ in $G'$. In this time, the set ${E'}_{M}$ of missing links should include the observed links in validation set $E_{P}$ as they were not used to training. In principle, the score of link in $E_{P}$ should be higher than that of link in ${E'}_{M}$ but not in $E_{P}$.

The average values of AUC and Precision [12] in 3-fold cross-validation were taken as evaluation indexes. For the *i*-th comparison in $N$ independent comparisons, we randomly selected two links from the set of missing links ${E'}_{M}$ (one of two should in the validation set, denoted as $l_{i}$ and the other should not existed in the validation set, denoted as $l_{j}$). If the score $S_{i}$ of link $l_{i}$ is higher than the score $S_{j}$ of link $l_{j}$, the prediction is considered to be unsatisfactory; If the score $S_{i}$ of link $l_{i}$ is lower than the score $S_{j}$ of link $l_{j}$, the prediction effect is considered to be satisfactory; If the score $S_{i}$ is equal to the score $S_{j}$, the prediction is random. Therefore, the AUC value was calculated as follows:

$$AUC=\frac{1}{N}\sum_{i=1}^{N} x_{i}，x_{i}=\left\{ \begin{aligned} 1, S_{i}>S_{j}; \\ 0.5, S_{i}=S_{j}; \\ 0, S_{i}<S_{j}. \end{aligned} \right.$$

where *N* is the number of independent comparisons, $x_{i}$ is the prediction effect of *i*-th comparison.

The evaluation metric Precision measures whether the top-L links order by the RWR score in set ${E'}_{M}$ is predicted accurately. In general, the links in $E_{P}$ should rank higher than the links in ${E'}_{M}$ but not in $E_{P}$. Precision is defined as the frequency of links in $E_{P}$ in $L$=100 random selection of ordered links:

$$Precision=\frac{1}{L}\sum_{i=1}^{L} x_{i}，x_{i}=\left\{ \begin{aligned} 1, l_{i}\in E_{P}; \\ 0, l_{i}\notin E_{P}. \end{aligned} \right.$$

Where $L$ is the number of random selection, $l_{i}$ represents the selected link in the *i*-th selection. Finally, considering both AUC and Precision, the optimal value of *r* is determined to be 0.61 (**Fig S11B**). After determining parameter in the RWR process, we could obtain the potential regulatory ability of the receptor on the downstream gene (TF).

***Determination of threshold for receptor-TF regulatory matrix based on cell line data***

We observed that in the regulatory potential matrix, most of the upstream receptors' regulatory potential to the downstream transcription factors is not zero. This is owing to that the receptor nodes can reach any protein nodes in the directed weighted graph via random walks except isolated nodes and nodes with zero in-degree. We therefore select a quantile cutoff θ (quantile.cutoff) to retain significant receptor-transcription factor interactions based on gene expression data of cell line perturbation experiments.

129 sets of cell line data (**Table S2**) were collected and analyzed to explore the regulation of upstream signal molecules (ligand, receptor) to downstream target genes via ligand treatment or receptor interference in a single cell line.

Firstly, three matrices were constructed according to the prior database: Ligand-receptor regulatory matrix $A$, receptor-transcription factor regulatory potential matrix $B$ and transcription factor-target gene regulatory matrix $C$. Denote $a_{i,j}$,$c_{k,t}$ and $b_{j,k}$to represent the number of pairing occurrences in the prior database (ligand-receptor pairing, transcription factor-target gene pairing) and the regulatory potential (receptor-transcription factor pairing), with $i, j, k, t$ indicating the index of ligand, receptor, transcription factor and target gene, respectively. For regulatory potential matrix $B$, we set the quantile cutoff $\theta$ to redefine $b_{j,k}$,:

$$b_{j,k}=\left\{ \begin{aligned} b_{j,k}, b_{j,k}\geq b_{\theta} \\ 0, b_{j,k}<b_{\theta} \end{aligned} \right.，\theta\in(0,1)$$

where $b_{\theta}$ is $\theta$-quantile threshold for the regulatory potential matrix $B$.

On the other hand, we analyzed differentially expressed genes (DEGs) using limma (padj<=0.05, |logfc| >= 1) for each cell line under specific perturbations (ligand treatment or receptor knockdown/mutant). The DEGs corresponding to a specific ligand or receptor was regarded as potential target genes regulated by a specific ligand ($L_{l}$) or receptor ($R_{r}$). Consequently, the ligand-target gene matrix $D_{l,m}$ and receptor-target gene matrix $E_{r,n}$ were defined as follows:

$$D=\left\{ \begin{aligned} 1, {TG}_{m} is the DEG of L_{l} \\ 0, {TG}_{m} is not the DEG of L_{l} \end{aligned} \right.$$

$$E=\left\{ \begin{aligned} 1, {TG}_{n} is the DEG of R_{r} \\ 0, {TG}_{n} is not the DEG of R_{r} \end{aligned} \right.$$

Define matrix${}^{1}{ABC}=sgn(A\cdot B\cdot C)$, and let $\tilde{F_{1}}$ be a submatrix of ${}^{1}\mathrm{ABC}$ that preserves the same column and row with $D$. Performing Hadamard product between $\tilde{F_{1}}$ and $D$ to get a ligand-target validation matrix $F1$ which integrates prior database information and cell line perturbation data information. Similarly, we can obtain matrix ${}^{1}{BC}$ and receptor-target gene validation matrix $F2$ based on the priori matrices, $C$ and $E$.

We selected optimal threshold by solving the following optimization problem,

$$\theta^{*}=\arg\max_{\theta} abs\left( \frac{\frac{\Sigma(F_{1})}{\#\left( F_{1} \right)}}{\frac{\Sigma(D)}{\#\left( D \right)}}+\frac{\frac{\Sigma(F_{2})}{\#\left( F_{2} \right)}}{\frac{\Sigma(E)}{\#\left( E \right)}}-\frac{\Sigma({}^{1}{ABC})}{\#\left( {}^{1}{ABC} \right)}-\frac{\Sigma({}^{1}{BC})}{\#\left( {}^{1}{BC} \right)} \right)$$

where $\#\left( \cdot\right)$ represents the total number of elements of the matrix, and $\Sigma\left( \cdot\right)$ represents the number of the nonzero elements of the matrix. The first two terms of the objective function were used to preserve the information of DEGs in the cell line data as much as possible, and the last two terms were used to ensure the sparsity of the regulatory matrix. A series of values for $\theta$ was screened to maximize the above objective function. Finally, the optimal value of cutoff $\theta$ was determined as 0.98 (**Fig S11C**).

**Text S3. Implementation of CytoTalk, NicheNet and MISTy**

Cytotalk, NicheNet and MISTy were selected for comparison with stMLnet since they could infer LR-Target gene link in inter- and intra-cellular signaling network based on transcriptomics data of single cells or capture spots. Here we describe the details of implementing these methods.

we used default gene filtering conditions (GeneFilterCutoff = 0.05 and BetaUpperLimit = 100) when running CytoTalk in Breast Cancer of ST data and glioma of scRNA-seq data. An undirected unweighted graph was constructed from the edge information from the optimal signal transduction networks analyzed by CytoTalk based on PCSF algorithm. The graph only kept ligands form sender and genes from receiver. The distance from the upstream regulatory molecule (ligand/receptor) to the downstream target gene was used as a regulation score for quantitative evaluation of AUCROC and AUCPR. For sender cells of different cell type, a specific target gene in the receiver cells might be affected by the same upstream regulatory molecule (ligand) but through multiple cell-type specific signal transduction networks. In this case, we integrated the optimal networks from specific-receiver and different sender and calculated the distance from all ligand to the target as the ultimate regulation score.

When implementing NicheNet, we used relatively loose gene filtering to obtain as many ligand-receptor pairs as possible for the ST data of breast cancer. The average expression of genes in the selected ligand-receptor pairs should be larger than 0.1. ICGs (i.e., interaction-changed genes) were inputted as target gene set of interest, and all activated ligands were used to evaluate the performance of prediction. For scRNA-seq data of glioma, the DEGs between resistant and sensitive groups were inputted as target gene set of interest. The ‘get_weighted_ligand_target_links’ function of NicheNet was used to obtain the activated ligand-target gene pairing and the weight was used as ligand’s regulatory ability score with respect to the downstream target genes. Different types of sender cells might interact with the receiver cells through the same ligand-receptor pair, but NicheNet constructed the intracellular signaling network based on the built-in database and did not distinguish between different sender cells. So we considered all activated ligands without distinguishing the sender cell types. In addition, NicheNet only inferred the ligand's regulatory ability to downstream target genes. For some receptors that were used for knockdown or mutation in the cell line perturbation data, we used their paired ligands for prediction of ligand/receptor-target regulation to evaluate the performance of NicheNet.

MISTy provides a way to study spatial relationships between genes based on the STdata from multiple views in terms of intra- and extra-cells. We employed MISTy with default parameters to predict the receptor-target regulation and ligand-target regulation by constructing intrinsic view and paraview, respectively. ICGs were used as targets of interest for MISTy prediction. MISTy calculated the importance scores of regulators (receptors or ligands) for the expression of each target from the specified view, which were then benchmarked with differential expressions of targets under specific ligand/receptor perturbations.

**Text S4. Simulation study**

To evaluate the quantitative model involved in stMLnet, we benchmarked stMLnet with a set of synthetic data of spatial gene expressions. We consider 5 ligands, 2 receptors, 3 TFs and 4 target genes. The ground truth of the multilayer network is shown in **Fig 2A**.

We simulate 3 types of cells, including 2 types of sender cells (SC_1_ and SC_2_) and 1 type of receiver cells (RC). These cells are randomly located at a fraction of grids within a 2-dimensional lattice (100$\times$100) that simulates a square domain of tissue slice ($\Omega\subset\boldsymbol{R}^{2}$). The ligand genes are expressed by the sender cells and the products are subject to diffusion. The receptor genes, TFs and TGs of the receiver cells do not diffuse across microenvironment.

The spatial-temporal changes of the extracellular ligands are modeled using the following reaction-diffusion equation

$$\frac{\partial[L_{i}]}{\partial t}=D_{i}\Delta\left[ L_{i} \right]+\sum_{k} r_{ik}\chi_{SC_{k}}(x)-d_{i}[L_{i}]$$

(S1)

where $[L_{i}]=[L_{i}](x,t)$ represents the concentration of the ligand $L_{i}$ at location $x\in\Omega$ and time $t$. $D_{i}$ and $d_{i}$ represents, respectively, the diffusion coefficient and degradation rate of the ligand $L_{i}$*.* $r_{ik}$ is the release rate of the ligand $L_{i}$ by sender cells $\mathrm{SC}_{k} (k=1 or 2)$. $\chi_{SC_{k}}(x)$ is an indicator function of the sender cells, taking value 1 where there is a sender cell at location $x$, and 0 otherwise. Random initial value and the no-flux boundary condition are imposed to the above equation.

The level of each receptor is assumed steady through the simulation. The activation of each TF within the receiver cells is modulated by the upstream LR signaling, which is described as follows,

$$\frac{\partial[TF_{l}]}{\partial t}=\sum_{j} \alpha_{jl}\sum_{i} b_{ij}[L_{i}]\cdot[R_{j}]-\beta_{l}[TF_{l}]$$

(S2)

where $[TF_{l}]=[TF_{l}](x,t)$ represents the activation level of the $TF_{l}$ at location $x$ and time $t$. $b_{ij}$ equals to 1 or 0, representing binding or non-binding between ligand $L_{i}$ and receptor $R_{j}$. $\alpha_{jl}$ is the activation coefficient of $TF_{l}$ by $R_{j}$. $\beta_{l}$ is the degradation rate of $TF_{l}$.

The expression of each target gene is regulated by TFs, which is described as follows,

$$\frac{\partial[TG_{s}]}{\partial t}=\sum_{l} \mu_{ls}[TF_{l}]-\gamma_{s}[TG_{s}]$$

(S3)

where $[TG_{s}]=[TG_{s}](x,t)$ represents the expression level of the target gene $TG_{s}$ at location $x$ and time $t$. $\mu_{ls}$ is the regulatory coefficient of $TG_{s}$ expression by $TF_{l}$. $\gamma_{s}$ is the degradation rate of $TG_{s}$.

To get the steady-state values of spatial gene expression, we solve the following equations:

$$-D_{i}\Delta\left[ L_{i} \right]=\sum_{k} r_{ik}\chi_{SC_{k}}\left( x \right)-d_{i}\left[ L_{i} \right]$$

$$\left. \frac{\partial[L_{i}]}{\partial\vec{n}} \right|_{\partial\Omega}=0$$

(S4)

$$[TF_{l}](x)=\frac{1}{\beta_{l}}\sum_{j} \alpha_{jl}\sum_{i} b_{ij}[L_{i}]\cdot[R_{j}]$$

(S5)

$$[TG_{s}](x)=\frac{1}{\gamma_{s}}\sum_{l} \mu_{ls}[TF_{l}]$$

(S6)

Equations (S4) were solved using finite difference method with five-point central difference scheme (see details below). The values of some parameters (e.g., $r_{ik}$, $\alpha_{jl}$, $\beta_{l}$, $\mu_{ls}$, $\gamma_{s}$ and $[R_{j}]$) in the above model were randomly sampled in each simulation. By simulating the above model with random parameter values 100 times, we got 100 sets of synthetic data each includes spatial expression values of 5 ligands, 2 receptors and 4 target genes and location coordinates of the SCs and RCs. **Fig S3C-H** illustrate a set of representative simulation data. Such spatial expression data was used as input of the random forest regression model in stMLnet (without using the prior information of the predefined multilayer network) to infer the regulation of TGs’ expression by LR pairs. The predicted importance scores for LR-TG regulations were benchmarked with the ground truth.

To numerically solve Equation (S4), we employed finite difference method with five-point central difference scheme. For simplicity, we wrote the Equation (S4) into the following form by ignoring index *i*:

| 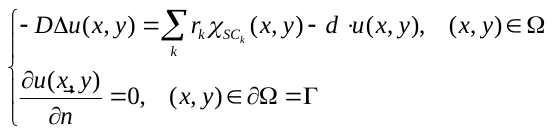, | (S7) |
| --- | --- |

where
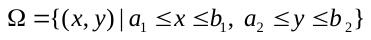
. We first uniformly divided the whole domain into
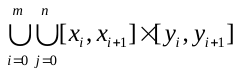
, here
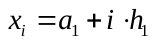
,
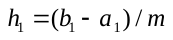
,
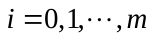
, and
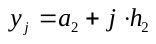
,
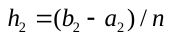
,
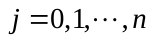
.

Assume that
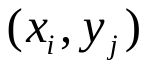
 is a regular interior point. Denote
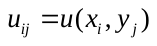
,
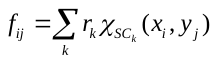
. By using five-point central difference stencils to approximate the diffusion term, we get

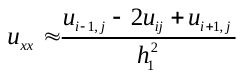
,

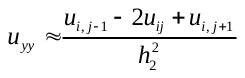
.

Therefore, at each interior point
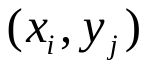
, we get the following difference equations

| 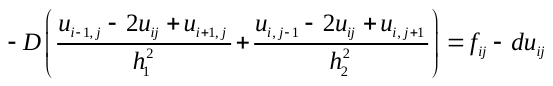,  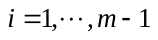, 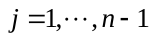. | (S8) |
| --- | --- |

Order the
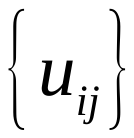
 and
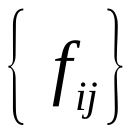
 into a 1-dimensional vector as follows

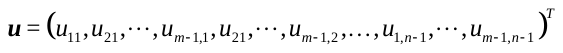
,

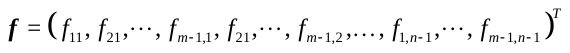
.

At the boundary points, by discretizing the boundary conditions, we have

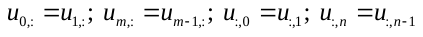
.

These constrains could be incorporated into the Equations (S8).

We next sought to represent the above Equations (S8) with matrix form.

Denote

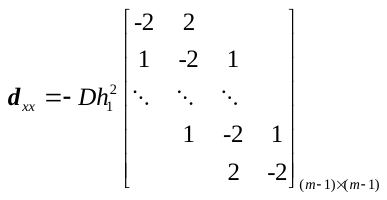
,

and

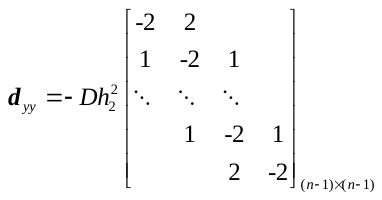
.

Then the discretization matrix of
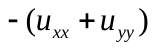
 could be represented as

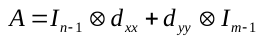
,

where
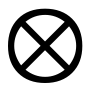
 stands for the Kronecker product, and
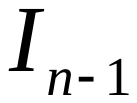
 for an identity matrix of order
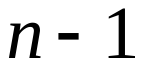
.

As such, the Equation (S8) could be written into the following matrix form

| 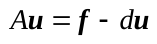, | (S9) |
| --- | --- |

Consequently,

| 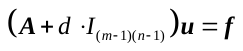. | (S10) |
| --- | --- |

Now the numerical solutions to the above linear algebraic equations (i.e., Equation (S10)) are readily solvable. In our simulation, we set
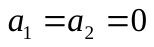
,
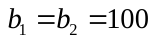
, and
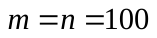
.

**Text S5. Comparison of prior databases**

We here discuss the difference of the prior databases in stMLnet and that in other tools such as NicheNet and Omnipath. stMLnet collected not only intercellular communication interactions (ligand-receptor interactions), but also intracellular signaling interactions (receptor-transcription factors, transcription factor-target gene interactions). At present, NicheNet collected interactions from 57 resources, while Omnipath collected interactions from more than 100 resources. The main differences between them and stMLnet are as follows.

(1) *Data collection resources and standards*. (i) At the ligand-receptor interaction level, stMLnet included information from NicheNet and Omnipath at the ligand-receptor interaction level, so its ligand-receptor pairs largely overlap with NicheNet and Omnipath (**Fig S11A**, left panel). (ii) At the level of intracellular signaling interactions, NicheNet collected not only a large amount of protein interaction information but also a lot of self-annotated interaction information. As such, signaling pairs in NicheNet were far more than that of stMLnet and Omnipath (**Fig S11A**, middle panel). (iii) At the transcriptional regulation level, stMLnet collected a large number of TF-TG interactions for better leveraging the expression data of cell line perturbation experiments to determine significant signaling path from specific receptors to TFs (**Text S2**). Therefore, stMLnet has much more information on regulatory interactions than NicheNet and Omnipath (**Fig S11A**, right panel).

(2) *Prediction of directed interactions*. stMLnet selected receptor-transcription factor interactions with highly regulatory potential by collecting information mainly from pathway databases (PathBank, SMPDB, Reactome, KEGG, BioCarta, NCI, PATHER, PharmGKB) (**Fig S11C**). While NichNet and Omnipath mainly collected such information from protein-protein interaction databases besides signaling pathway databases. We know that receptor-TF signal transduction has obvious direction. While the information collected from pathway database generally has obvious signal transmission direction (ligand-receptor-downstream signal molecule-transcription factor-target gene), but there are a lot of unknown direction of interaction in protein-protein interaction database. Therefore, bias may be introduced by using protein-protein interaction database information when constructing a directed graph to predict receptor-transcriptional factor interactions. Therefore, NichNet and Omnipath, although collecting more abundant data resources, were not as effective as stMLnet in predicting directed interactions (**Fig S11B**).

**Text S6. Data collection and processing**

***Single-cell resolution ST datasets***

Several single-cell resolution ST datasets generated from different technologies, including seqFISH+, Slide-seq v2 and MERFISH, were collected for demonstration the applicability of stMLnet. The seqFISH+ data of mouse cortex contains 10000 genes measured in 913 cells [13]. The Slide-seq v2 data of mouse hippocampus contains 53173 cells and 13854 genes [14]. The MERFISH data is a 3D spatial expression dataset of mouse hypothalamus preoptic region, consisting of 155 genes from ~1 million single cells in 12 non-continuous tissue slices [15]. These datasets were downloaded, preprocessed and annotated by using Seurat and Giotto according to the procedures described in the original tutorial. The processed dataset was used as input of stMLnet. The unannotated cells were filtered. The ligand and receptor genes were selected by their mean gene expression and gene expression percentage. For the seqFISH+ dataset and Slide-seq v2 dataset, cell type-specific genes selected by using ‘findMarkers’ function in Seurat were used as the target gene set of interest for input. For the MERFISH dataset, cell type-specific genes selected via “findMarkers_one_vs_all” function in Giotto were used as the target gene set of interest for input of stMLnet.

***Breast cancer dataset***

The spatial transcriptomics (ST) data of breast cancer was obtained from the 10x Genomics website (https://www.10xgenomics.com/resources/datasets, Visium Demonstration, Human Breast Cancer, Block A Section 1). The preprocessing of the ST data, including quality control, normalization, and dimension reduction (tSNE analysis), was conducted using standard pipeline of Seurat. The cell type annotation information in a scRNA-seq dataset (GSE118389) [16] was mapped to each spot of the ST data (‘FindTransferAnchors’ and ‘TransferData’ in Seurat v3.2.3 [17]). Furthermore, the cell type-specific gene expression for the dominant cell type in each spot was calculated using ‘get_decomposed_data’ function in RCTD v1.1.0 [18] with the aid of the cell type proportion matrix from Seurat output and the above scRNA-seq reference dataset. We also employed Giotto [19] to get ICGs by using ‘findICG’ function and ‘filterICG’ function with default parameters, which were used as the input target gene set of interest for stMLnet. The ST data was imputed using Seurat ‘runALRA’ function to mitigate the sparsity of the expression matrix for the LR signaling quantification and random forest regression.

***Glioma dataset***

The ST data of glioma was generated by 10X Visium technology [20]. This dataset includes spatially resolved transcriptomics data of 28 samples. In this study, we selected sample ‘#UKF304_T’ for subsequent analysis. The preprocessing of the ST data, including quality control, normalization, and dimension reduction (tSNE analysis), was conducted using standard pipeline of Seurat. Following the original study [20], the SPOTlight method was used to decompose each spot into individual cell type with the aid of scRNA-seq data (GSE84465). Furthermore, we used the Seurat 'runALRA' function to perform imputation on the expression matrix of ST data to reduce the impact of gene expression sparsity on subsequent analyses. And then, we adopted the ‘get_decomposed_data’ function in RCTD v1.1.0 to calculate the cell type-specific gene expression for the dominant cell type in each spot.

To select target genes of interest as input for stMLnet, we collected a set of RNA-seq data of the isolated glioma cells or macrophages from three groups of mice with different responses to the CSF1R inhibitor treatment (Reb, rebound; EP, endpoint; Vel, vehicle) (GSE69104 [21]). After CPM standardization, low-expression genes with expression levels below 0.5 in more than 6 samples were filtered. We calculated differentially expressed genes (DEGs) in macrophages between different groups (Reb v.s. EP, EP v.s. Veh) using limma test (|logfc|>1, p.adj<0.1). The DEGs between Reb and EP could be viewed as resistance-related genes, which were input to stMLnet as potential target genes in macrophages or tumor cells to infer their upstream regulators.

For other cell types (e.g., T cells and oligodendrocytes), we used highly expressed genes in these cell types, analyzed from the scRNA-seq data by using ‘FindMarkers’ function in Seurat (|logfc|>2, p.adj<0.05, pct>=0.1), as respective target genes of interest for stMLnet input.

In addition, the DEGs in macrophages between EP and Veh groups could be viewed as genes responsive to CSF1R inhibition and thus potentially regulated by CSF1R, which were used to test the prediction of stMLnet with respect to the CSF1R-regulated target genes.

***COVID-19 dataset***

The ST data of COVID-19-infected human lung tissue was processed with the original code deposited at Mendeley: <https://doi.org/10.17632/xjtv62ncwr.1>. Following the original study [22], the NNMF method and the Pearson correlation were used to calculate spot-factor matrix and factor-celltype matrix to calculate cell type proportion for each spot in the ST data. More specifically, we firstly merged the similar subtypes according to the reference scRNA-seq dataset [23] and then two truncation parameters (factor_cutoff = 0.5 and celltype_cutoff = 0.5) related to spot-factor matrix and factor-celltype matrix were utilized to avoid excessive dispersion of cell type proportion. The cell type proportion matrix was defined as the Hadamard product of the above two matrices. The cell type-specific gene expression for the first two dominant cell types in each spot was calculated using ‘get_decomposed_data’ function in RCTD v1.1.0 [18]. The cell type-specific DEGs in the ST data in each type of receiver cells (analyzed using ‘FindMarkers’ function in Seurat) were selected as the target genes of interest for input of stMLnet to infer multilayer signaling networks.

***Cell lines data***

To calibrate parameters for prior information integration, we collected 129 datasets of cell line gene expression that were treated with ligands or perturbed by receptor knockout/mutation from the GEO database (**Table S6**), involving 23 tissues, 65 cell lines, 42 ligands and 18 receptors. We used limma [24] to select DEGs of each cell line (before vs. after perturbation) under criteria of |logfc|>1 and p.adj<0.05. Ultimately 70 datasets, each with more than 50 DEGs, were used for correction of prior databases in this study.

**Figure S1**

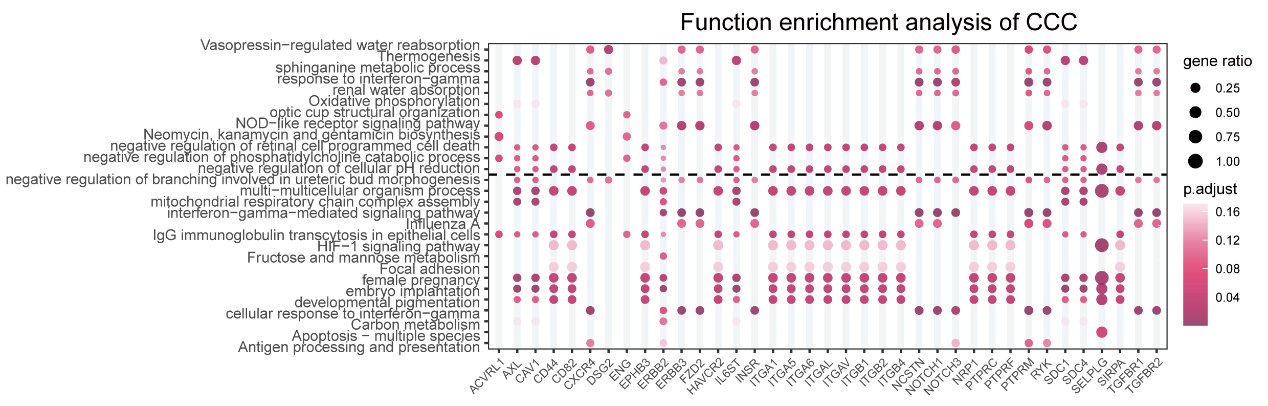

**Figure S1.** **Enrichment analysis for the multilayer signaling networks of breast cancer**. Heatmap of functional enrichment for CCC. The GO biological process (BP) (up) or KEGG (below) enrichment of the downstream targets for a set of upstream receptors during CCI with T cells as senders and malignant cells as receivers is shown. Top-ranked function terms according to p-values were prioritized for visualization.

**Figure S2**

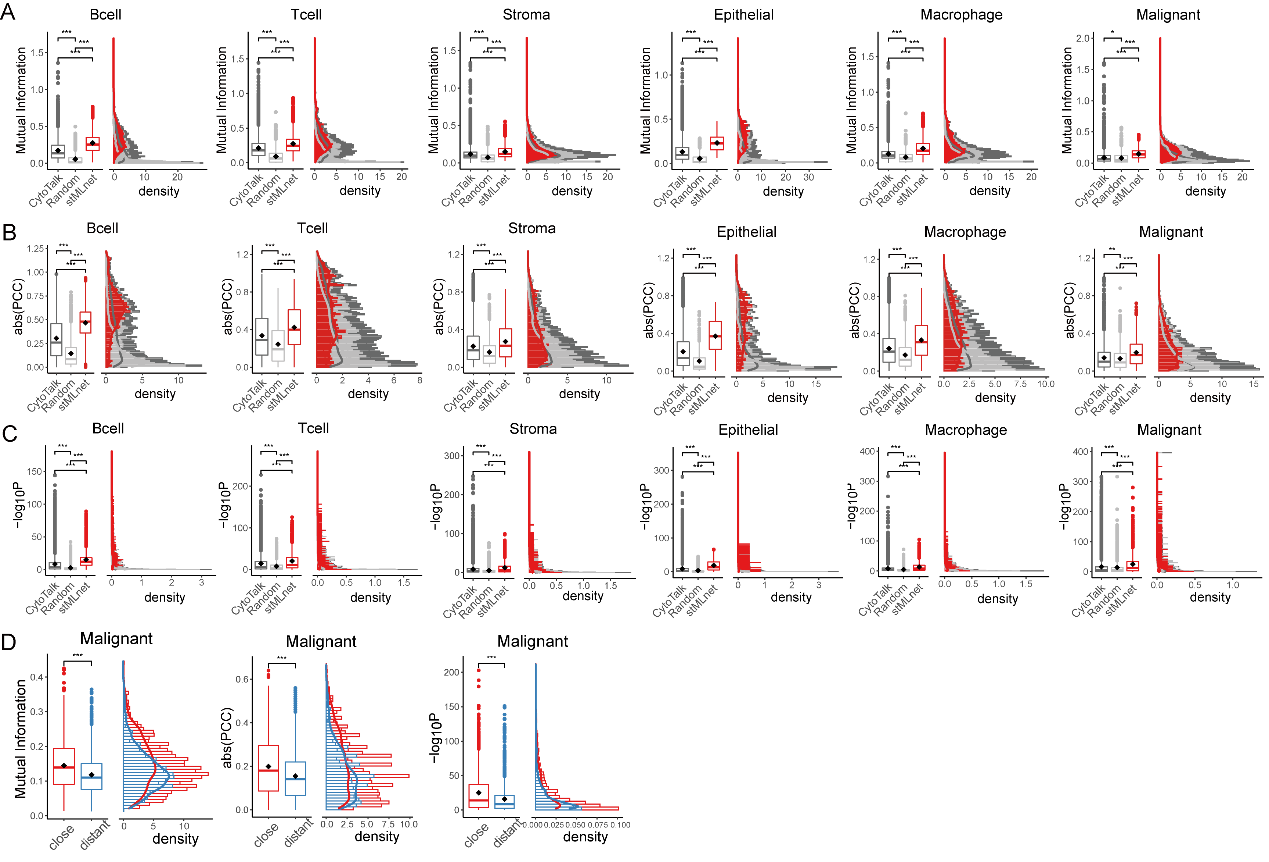

**Figure S2**. **Verifying stMLnet based on LR-target correlations**. (**A**) Distributions of mutual information between LR signaling activities and target gene expressions (${\mathrm{LR}_{k_{t}}\sim TG}_{t}$) in different cell types. The LR-target pairings predicted by stMLnet had larger mutual information values than other methods (e.g., CytoTalk) and random pairing. (**B**) Distributions of absolute values of PCC (abs(PCC)) between LR signaling activities and target gene expressions in different cell types. The LR-targets pairings predicted by stMLnet possessed larger abs(PCC) values than other methods (e.g., CytoTalk) and random pairing. (**C**) Distributions of –log10(p value) of the PCC between LR signaling activities and target gene expressions in different cell types. The LR-targets pairings predicted by stMLnet possessed larger values of –log10(p value of PCC) than other methods (e.g., CytoTalk) and random pairing. (**D**) Correlation of ${\mathrm{LR}_{k_{t}}\sim TG}_{t}$ predicted by stMLnet in malignant cells stratified by close or far distance to the sender cells. The close group had higher correlation than the distant group, assessed by the mutual information, the absolute value of PCC, and the –log10(p value of PCC).

**Figure S3**

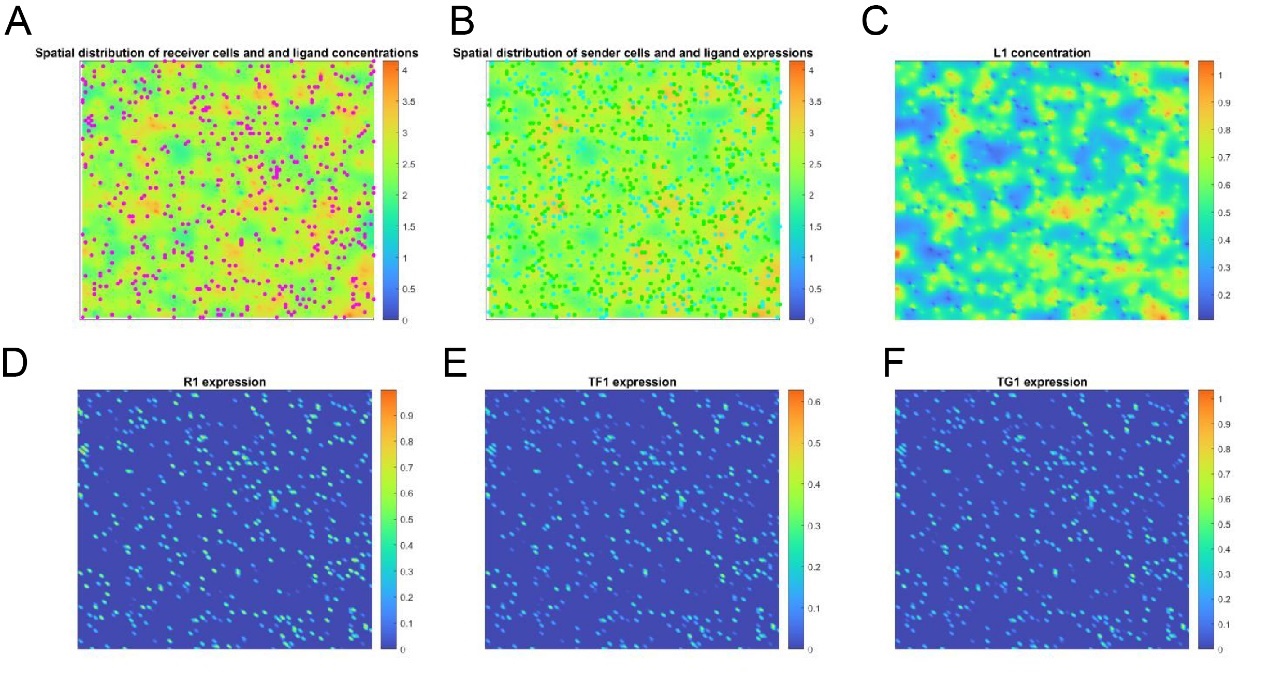

**Figure S3.** **Illustration for the simulation study**. (**A-F**) A representative simulation realized with a set of randomly sampled parameters. Shown are spatial distributions of receiver cells (**A**) and sender cells (**B**) with total ligand concentration as background, L1 concentration (**C**), R1 expression (**D**), TF1 activation (**E**) and TG1 expression (**F**), respectively.

**Figure S4**

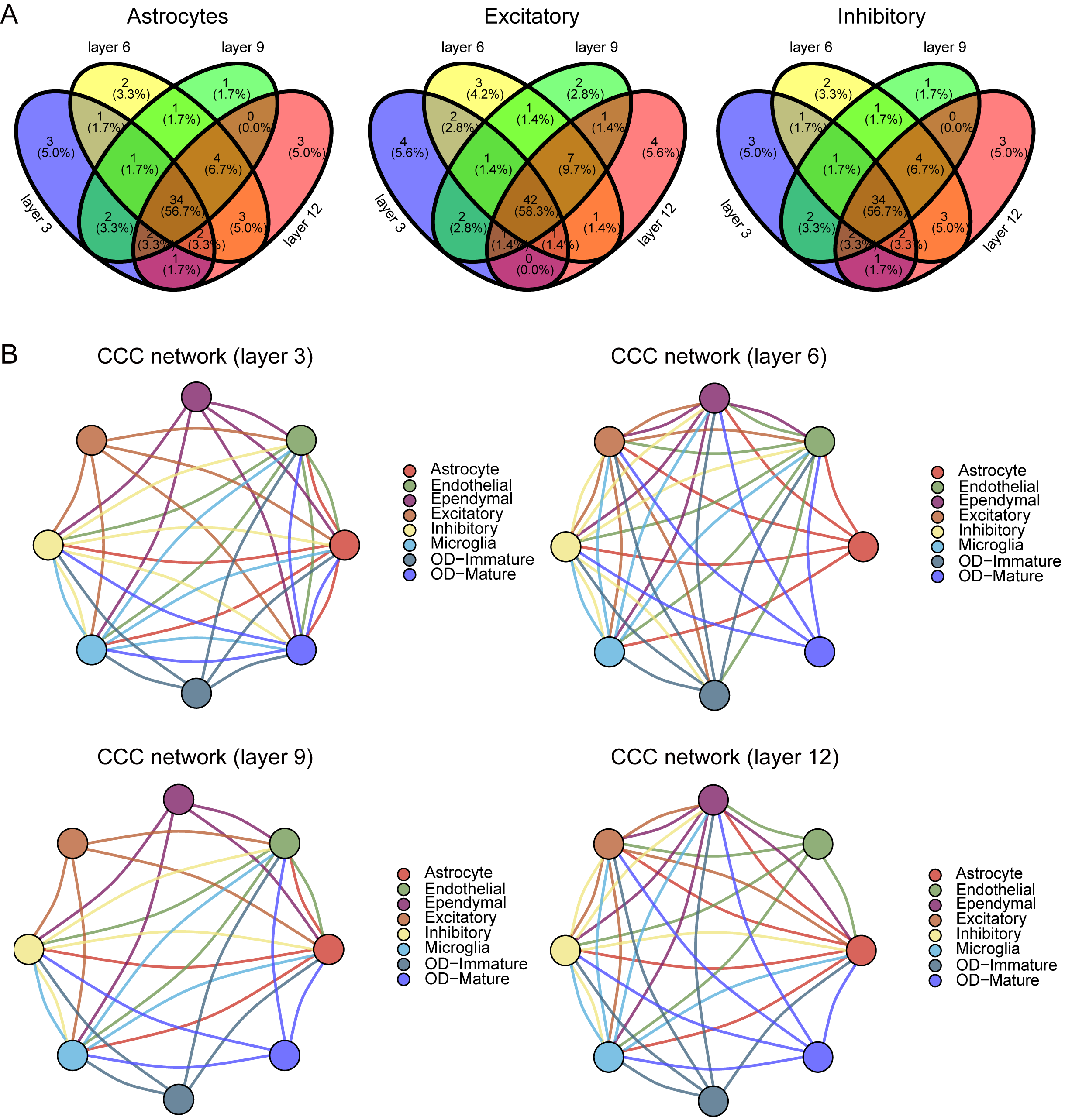

**Figure S4. Comparison of cell communication in different layers of MERFISH data.** (**A**) The potential target genes in different layers of MERFISH data. (**B**) The CCC network of mouse brain's preoptic regions in different layers. The width of the edge represents the number of target genes in the receiver cell type.

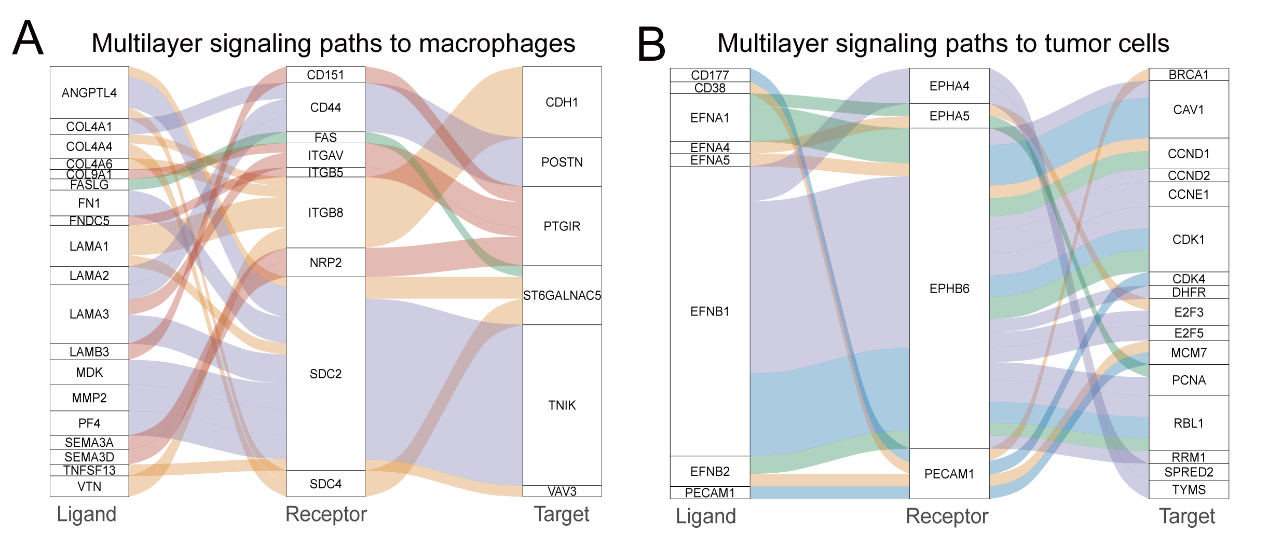
**Figure S5**

**Figure S5. The waterfall plot of the multilayer signaling networks in the Glioma ST dataset.** (**A**) The multilayer signaling paths from upstream LR pairs (top-ranked) to their downstream targets in macrophages. (**B**) The multilayer signaling paths from upstream LR pairs (top-ranked) to their downstream targets in tumor cells. Different colors of paths represent cellular sources (sender cells) of the ligand signaling, and the width of the path represents the importance score for each LR~TG regulation.

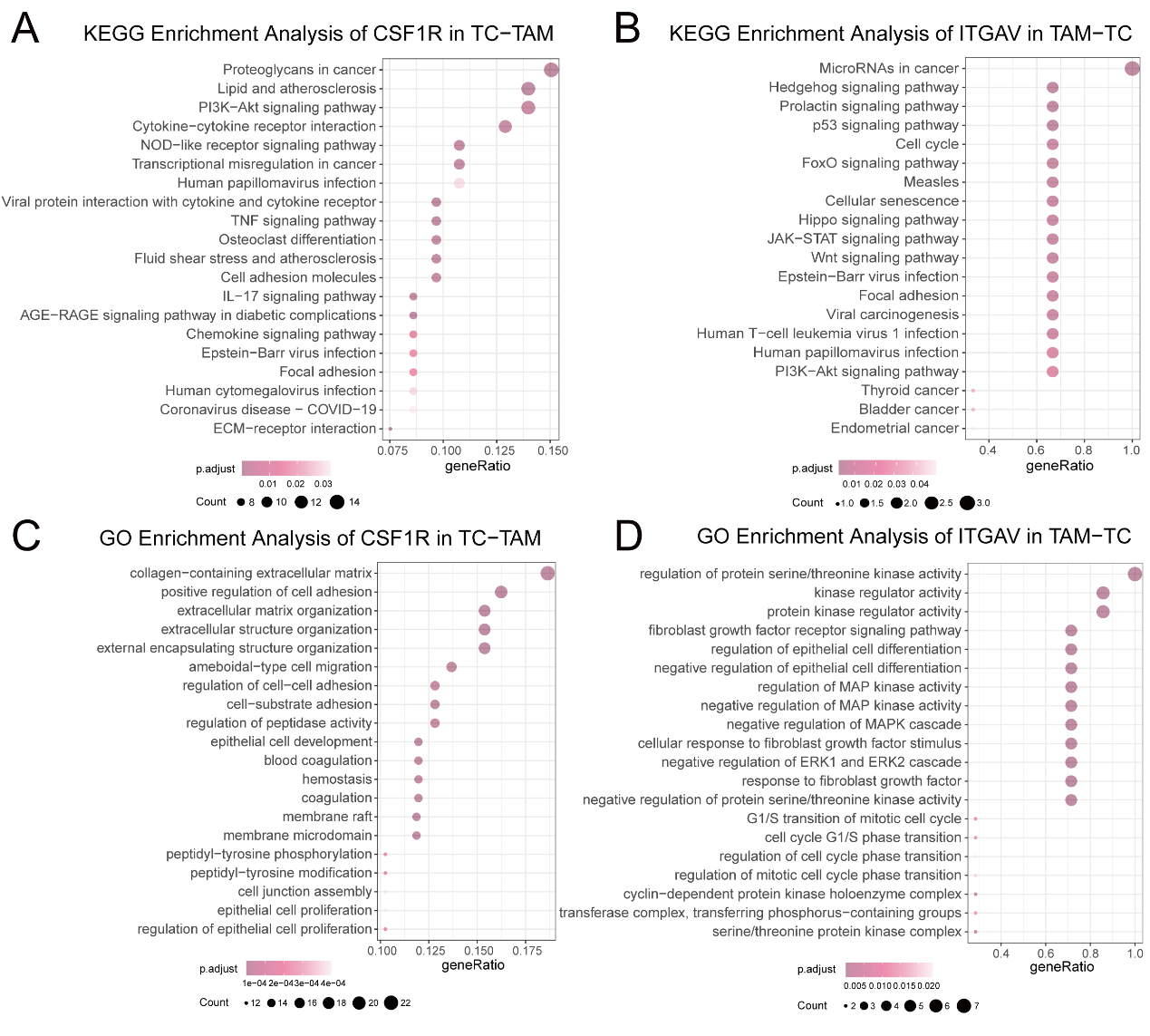
**Figure S6**

**Figure S6**. **Enrichment analysis for the multilayer signaling networks of glioma**. (**A**) The KEGG enrichment analysis of downstream targets of CSF1R in macrophages regulated by malignant cells. (**B**) The KEGG enrichment analysis of downstream targets of ITGAV in malignant cells regulated by macrophages. (**C**) The GO enrichment analysis of downstream targets of CSF1R in macrophages regulated by malignant cells. (**D**) The GO enrichment analysis of downstream targets of ITGAV in malignant cells regulated by macrophages.

**Figure S7**

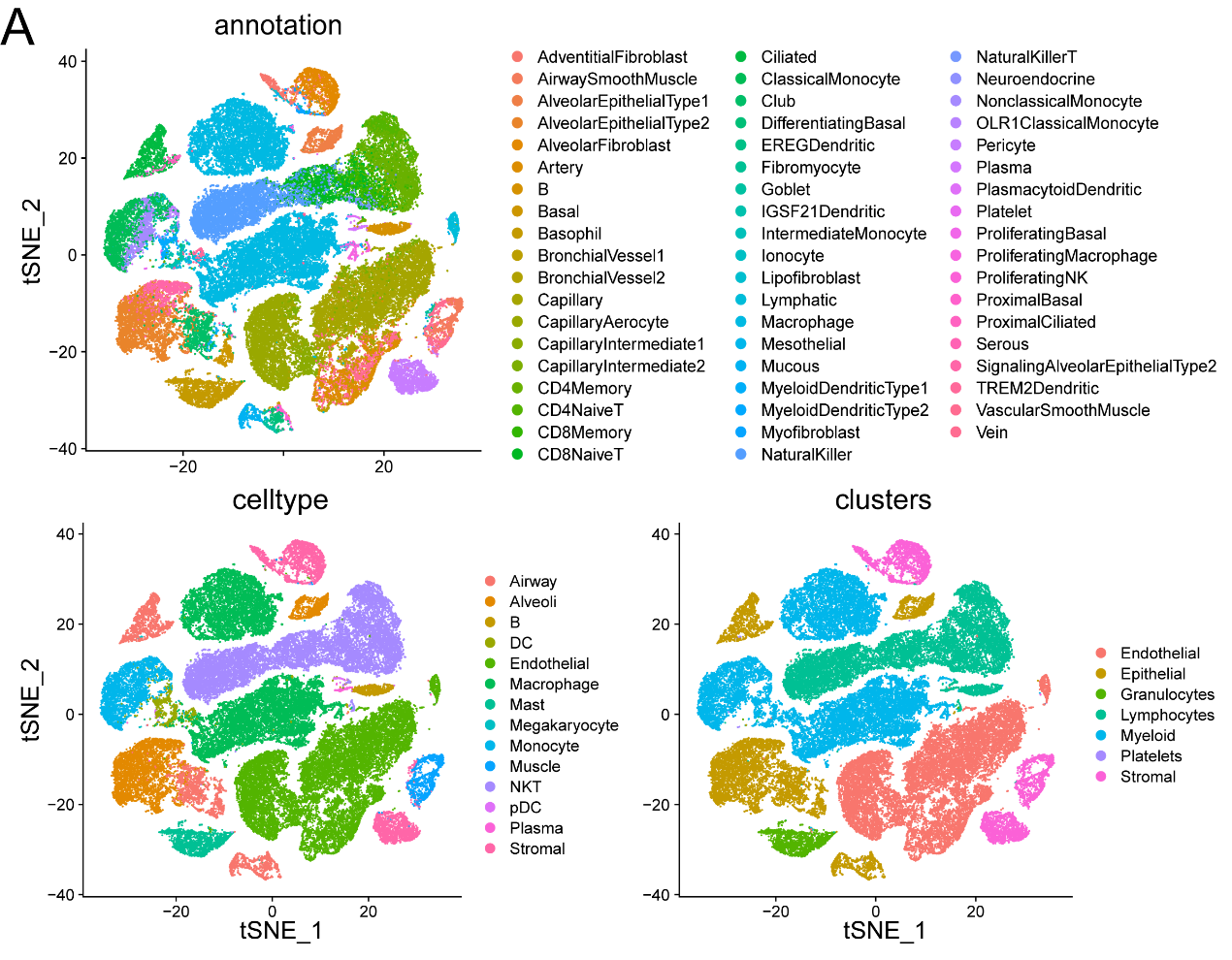

**Figure S7.** **The cell type annotation for the scRNA-seq data of COVID-19**. Shown are TSNE of expression profile of scRNA-seq data from views of annotations, cell types and clusters.

**Figure S8**

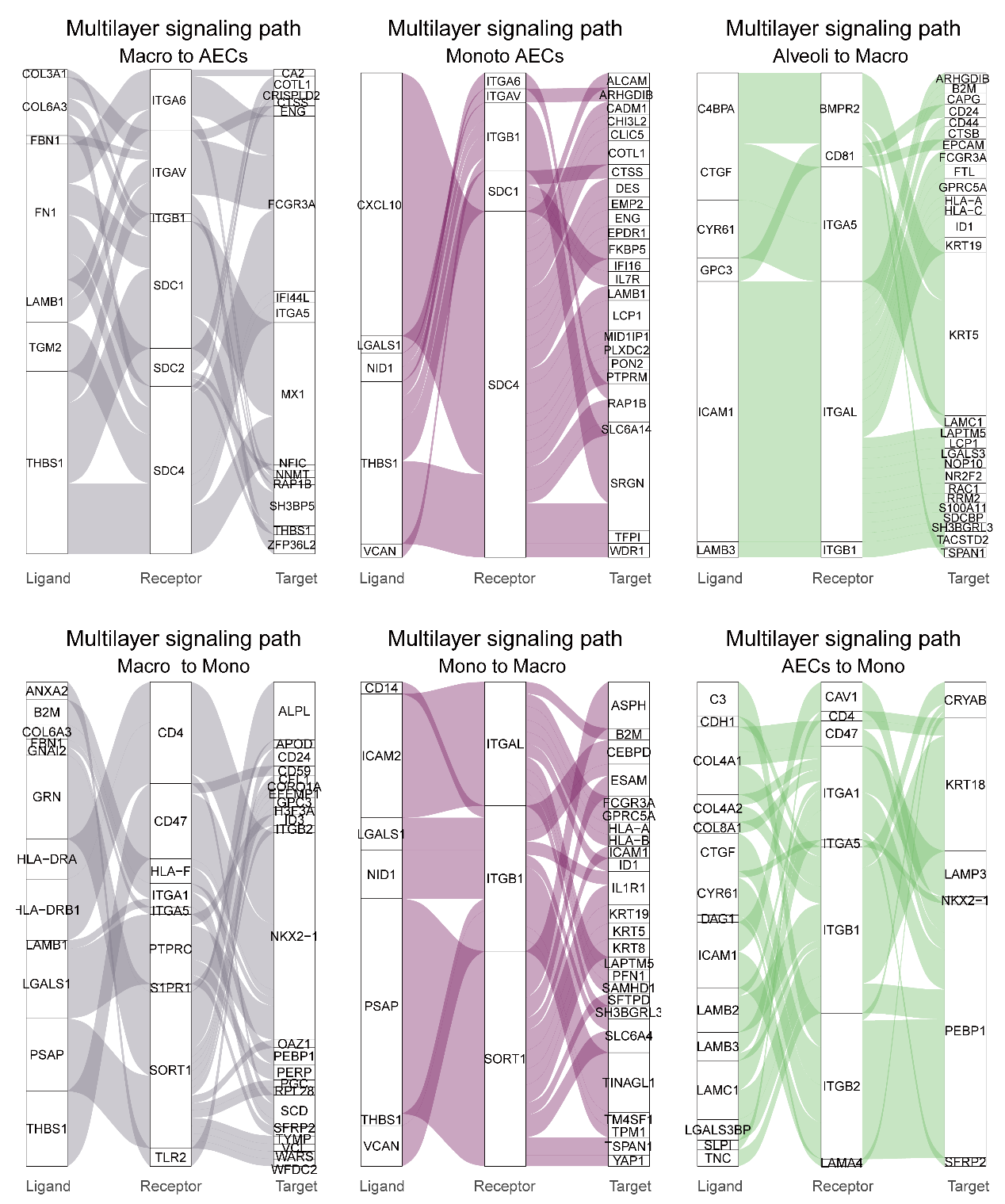

**Figure S8**. **The waterfall plot of the multilayer signaling network for the cellular feedback circuits inferred from the COVID-19 ST dataset.** Shown are regulatory paths from upstream LR pairs (top ranked) to their downstream targets for each pair of cell types among AECs, macrophages, and monocytes. Different colors of paths represent cellular source (sender cells) of the ligand signaling, and width of the path represents the importance score of each LR~TG regulation.

**Figure S9**

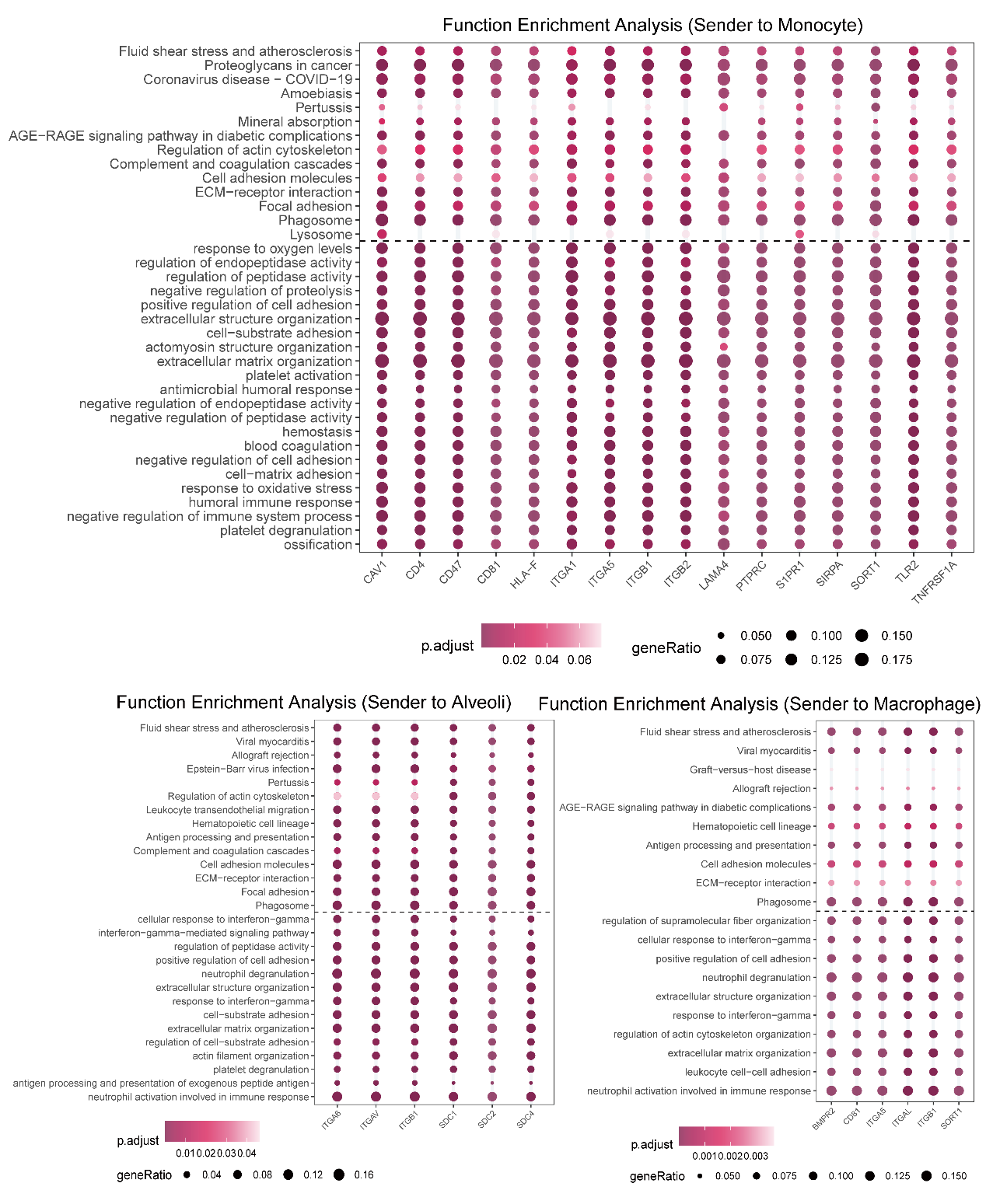

Figure S9. Heatmap of functional enrichment for cellular feedback circuits inferred from the COVID-19 ST dataset. Shown are GO biological process (up) or KEGG (below) enrichment results of the intracellular target genes downstream of the upstream receptors involved in the multilayer networks for feedback circuits among AECs, macrophages and monocytes. Top ranked gene terms according to p values were prioritized for visualization.

**Figure S10**

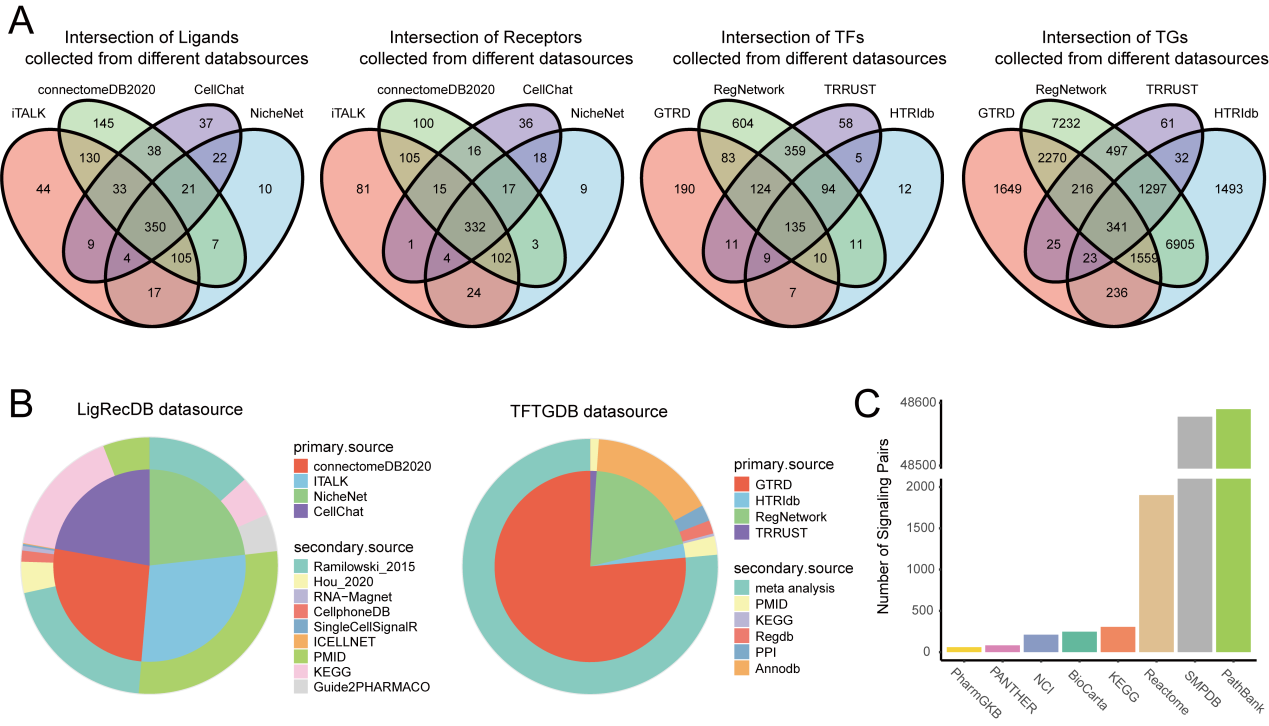

**Figure S10**. **The collection and sorting of prior knowledge database**. (**A**) The distribution of different types of molecules (ligands, receptors, TFs and target genes) recorded in LigRecDB and TFTGDB prior databases. (**B**) The distribution of different types of interactions (LR pairs or TF-TG pairs) collected in LigRecDB and TFTGDB prior databases. (**C**) Numbers of molecular interactions within signaling pathways collected in different pathway databases.

**Figure S11**

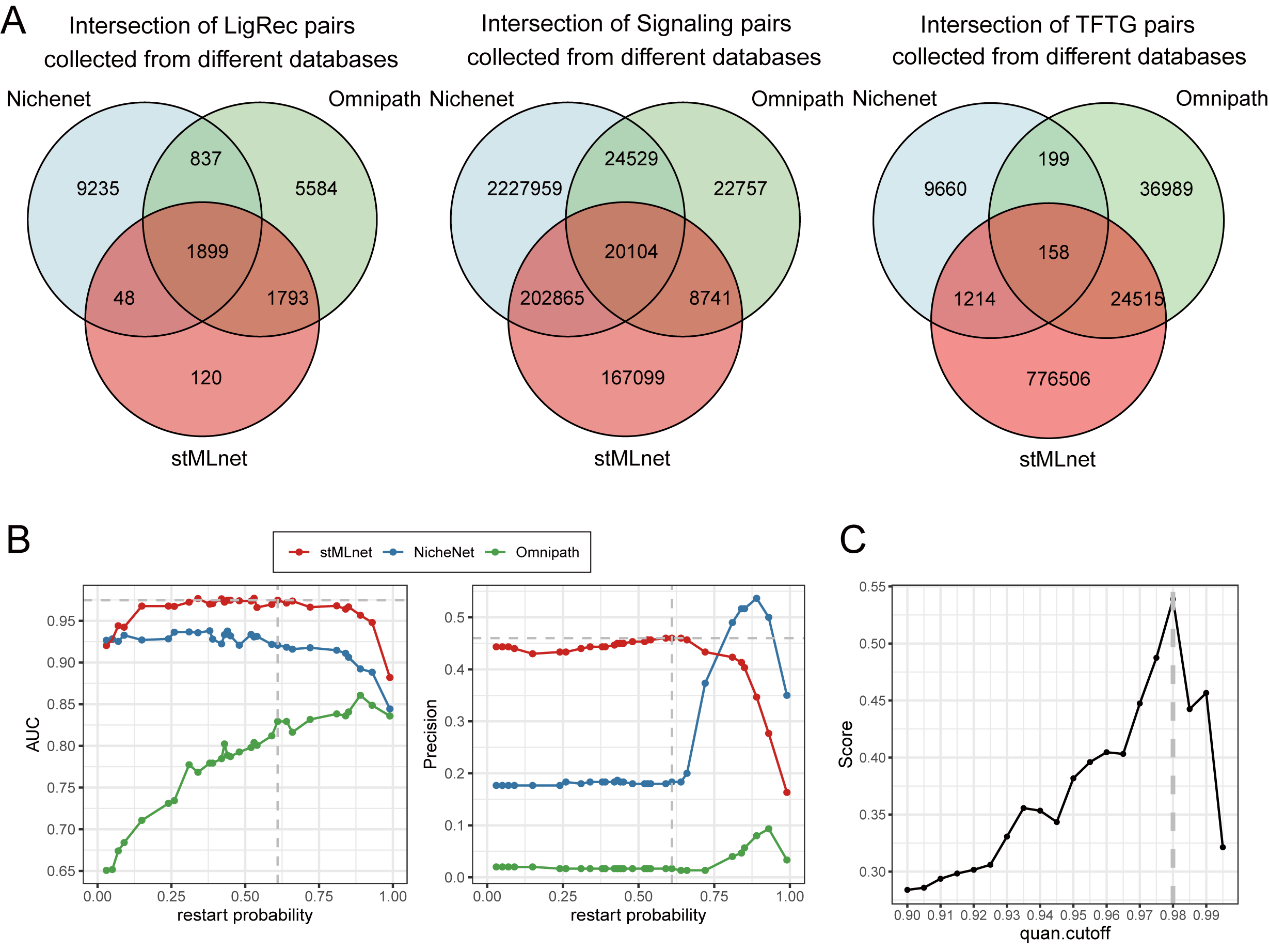

**Figure S11**. **Comparison and parameterization of prior knowledge databases**. (**A**) comparison of interaction pairs (LR, signaling pairs and TF-TG) in stMLnet, NicheNet and CytoTalk. (**B**) The determination of value of restart parameter of the random walk algorithm based on AUC and Precision. stMLnet was compared with NicheNet and Omnipath. (**C**) The determination of percentile cutoff value (quan.cutoff) for RecTF matrix derived from random walk algorithm using 74 datasets of gene expression in 48 cell lines with various stimulations or perturbations. The values in RecTF matrix represent predicted probabilities of interactions between the upstream and downstream signaling molecules. If the RecTF value was changed to 0 if it was less than quan.cutoff. Score in the y axis is an index defined taking into account both the sparsity of RecTF matrix and the information in the cell line datasets. The quan.cutoff was optimized to maximize the Score.
